# Large-scale Genetic Analysis Identifies 66 Novel Loci for Asthma

**DOI:** 10.1101/749598

**Authors:** Yi Han, Qiong Jia, Pedram Shafiei Jahani, Benjamin P. Hurrell, Calvin Pan, Pin Huang, Janet Gukasyan, Nicholas C. Woodward, Eleazar Eskin, Frank Gilliland, Omid Akbari, Jaana A. Hartiala, Hooman Allayee

**Author notes:** Address correspondence and reprint requests to: Hooman Allayee, PhD, Department of Preventive Medicine, Keck School of Medicine of USC, 2250 Alcazar Street, CSC202, Los Angeles, CA 90033. These authors contributed equally to this work (Jaana A. Hartiala, Hooman Allayee).

## Abstract

We carried out a genome-wide association study (GWAS) for asthma in UK Biobank, followed by a meta-analysis with results from the Trans-National Asthma Genetic Consortium (TAGC). 66 novel genomic regions were identified, bringing the number of known asthma susceptibility loci to 211. Significant gene-sex interactions were also observed where susceptibility alleles, either individually or as a function of polygenic risk scores, increased asthma risk to a greater extent in men than women. Bioinformatics analyses demonstrated that asthma-associated variants were enriched for colocalizing to regions of open chromatic in immune cells and identified candidate causal genes at 52 of the novel loci, including *CD52*. An anti-CD52 (α-CD52) antibody mimicked the immune cell-depleting effects of an FDA-approved human α-CD52 antibody and reduced allergen-induced airway hyperreactivity in mice. These results further elucidate the genetic architecture of asthma, provide evidence that the immune system plays a prominent role in its pathogenesis, and suggest that *CD52* represents a potentially novel therapeutic target for treating asthma.

## Introduction

Asthma is a global chronic respiratory disease affecting over 300 million people and a significant cause of premature death and reduced quality of life^1^. The disease develops through repeated cycles of inflammation, often in response to environmental triggers, that lead to progressive obstruction of the airways and deterioration of respiratory function^2^. While advances have been made in understanding these complex pathogenic processes, translation of research findings to development of novel therapies has not progressed rapidly enough to address the unmet clinical needs of many patients, particularly those with severe forms of asthma.

It is generally accepted that susceptibility to asthma is a combination of genetic predisposition, exposure to environmental factors, and their interactions. Consistent with this notion, estimates for the heritability of asthma have ranged between 35%-95%^3^, although those based on twin studies have been lower^4^. Genome-wide association studies (GWAS) have confirmed the contribution of genetic factors to risk of asthma with over 140 susceptibility loci having been identified over the last decade^5–24^. However, the risk alleles, most of which are common in the population, still only explain a small fraction of the overall heritability for asthma^25,26^. This observation implies either the existence of additional variants with smaller effect sizes, rare susceptibility alleles, and/or interactions between genes and environmental factors. In this regard, rare variants do not appear to explain a significant proportion of the “missing heritability” for asthma^27^. Gene-environment (GxE) interactions also remain poorly understood due to the inherent difficulties of carrying out such studies in humans, particularly with respect to accurate exposure assessment, adequately powered sample sizes with both genetic and exposure data, and the heterogeneous nature of asthma itself^28,29^. Thus, additional studies are needed to further elucidate the genetic architecture of asthma and determine the applicability of the findings for translational purposes.

Here, we identified 66 novel risk loci for asthma through a large-scale genetic study with 88,486 cases and 447,859 controls from UK Biobank and the Trans-National Asthma Genetic Consortium (TAGC). Follow up analyses demonstrated interactions between asthma susceptibility alleles and sex, provide further evidence that the immune system plays a prominent role in asthma pathogenesis, and functionally validated *CD52* as one candidate causal gene with the potential of being a novel therapeutic target.

## Results

### GWAS in UK Biobank

To further elucidate the genetic architecture of asthma, we first carried out a genome-wide association study (GWAS) in 64,538 cases and 329,321 controls from the UK Biobank (**Fig. 1** and **Supplemental Table 1**). This GWAS analysis identified 32,813 significantly associated SNPs distributed among 145 loci (**Supplemental Fig. 1** and **Supplemental Table 2**). Of these regions, 41 loci were novel and not previously known to be associated with asthma or the combined phenotype of asthma and allergic disease. The remaining 104 overlapped with the 145 loci previously identified for either asthma alone in adults, children, or subjects from multiple ancestries, or the combined phenotype of asthma, eczema, and hay fever^5–24^ (**Supplemental Table 2**).

**Figure 1.**
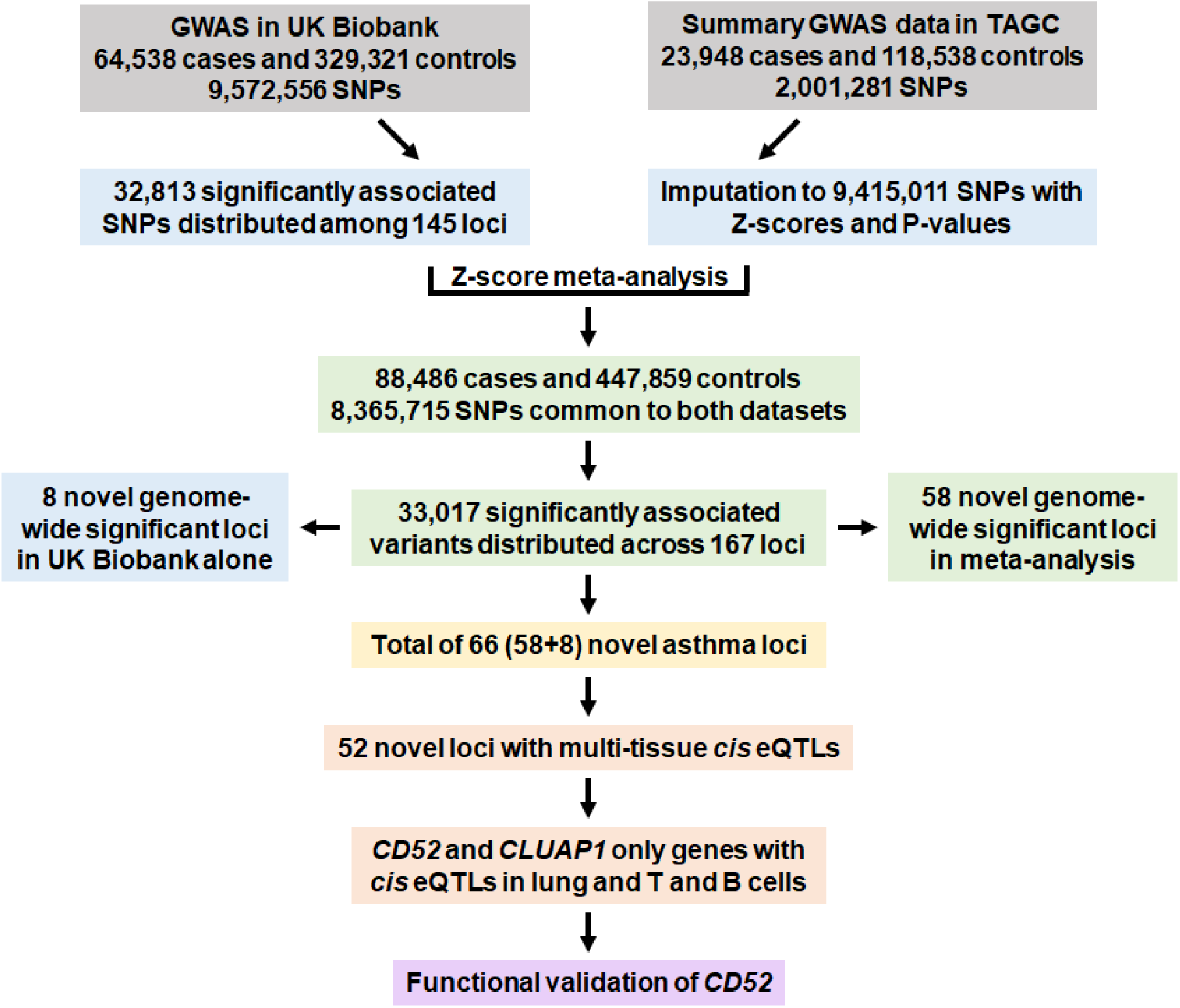
Overview of genetic and functional analyses. A GWAS was first carried out using primary level data in UK Biobank with ∼9.5 million SNPs. In parallel, summary statistics were imputed based on publicly available GWAS data from TAGC. The results were combined in a Z-score meta-analysis that included 88,484 asthma cases and 447,859 controls with 8,365,715 variants common to both datasets. 58 previously unknown genome-wide significant loci for asthma were identified in the meta-analysis. Combined with the eight novel genome-wide significant loci from the GWAS analysis in UK Biobank alone, a total of 66 novel asthma susceptibility loci were identified. Follow up bioinformatics and eQTL analyses prioritized *CD52* for *in vivo* functional validation studies.

### Meta-analyses of GWAS Data for Asthma in UK Biobank and TAGC

We next meta-analyzed the GWAS results from UK Biobank with publicly available summary data from the TAGC^21,30^ (**Fig. 1**). After harmonizing the summary GWAS results from TAGC to match the data in UK Biobank (see Methods for additional details), we performed a Z-score meta-analysis for asthma with 8,365,715 SNPs common to both datasets and a total of 88,486 cases and 447,859 controls (**Supplemental Table 1**). The meta-analysis revealed 33,017 significantly associated variants distributed across 167 loci (**Fig. 2; Supplemental Fig. 2; Supplemental Table 3**). Of these, 58 loci were novel and identified for asthma for the first time herein (**Fig. 2** and **Table 1**) whereas the other 109 overlapped with the 145 previously known loci (**Supplemental Table 3**).

**Figure 2.**
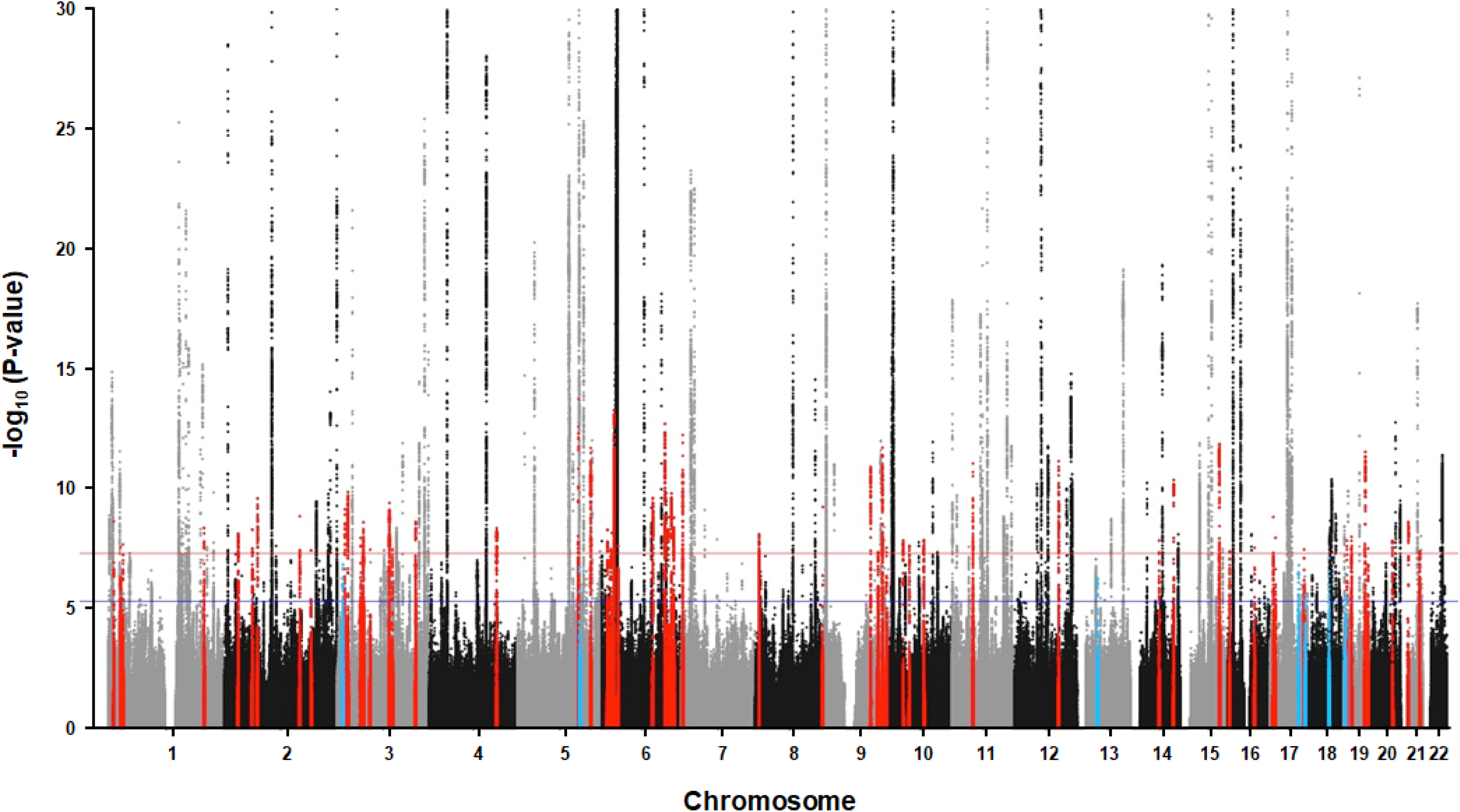
GWAS analyses in UK Biobank and TAGC identifies 66 novel loci for asthma. Of the 66 novel loci, 58 were significantly associated with asthma in the meta-analysis with UK Biobank and TAGC (red dots). Eight loci were associated with asthma in the GWAS analysis with UK Biobank alone but fell slightly below the genome-wide significance threshold in the meta-analysis with TAGC (light blue dots). The Z-score meta-analysis included a total of 88,486 cases and 447,859 controls from UK Biobank (64,538 asthma cases and 329,321 controls) and TAGC (23,948 asthma cases and 118,538 controls) and 8,365,715 SNPs common to both datasets. Genome-wide thresholds for significant (P=5.0×10^-8^) and suggestive (P=5.0×10^-6^) association are indicated by the horizontal red and dark blue lines, respectively. P-values are truncated at −log_10_(P)=30.

**Table 1.**
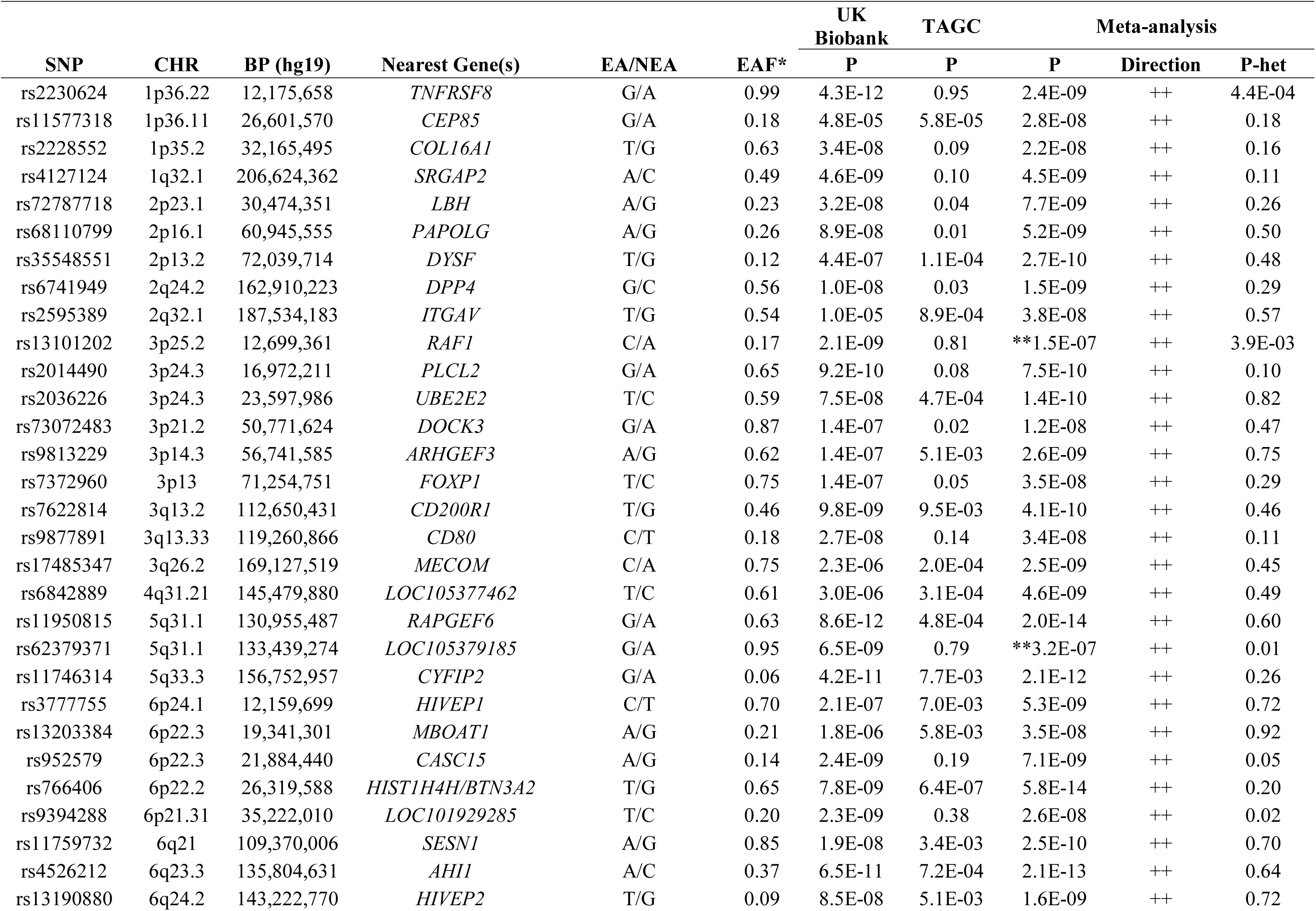
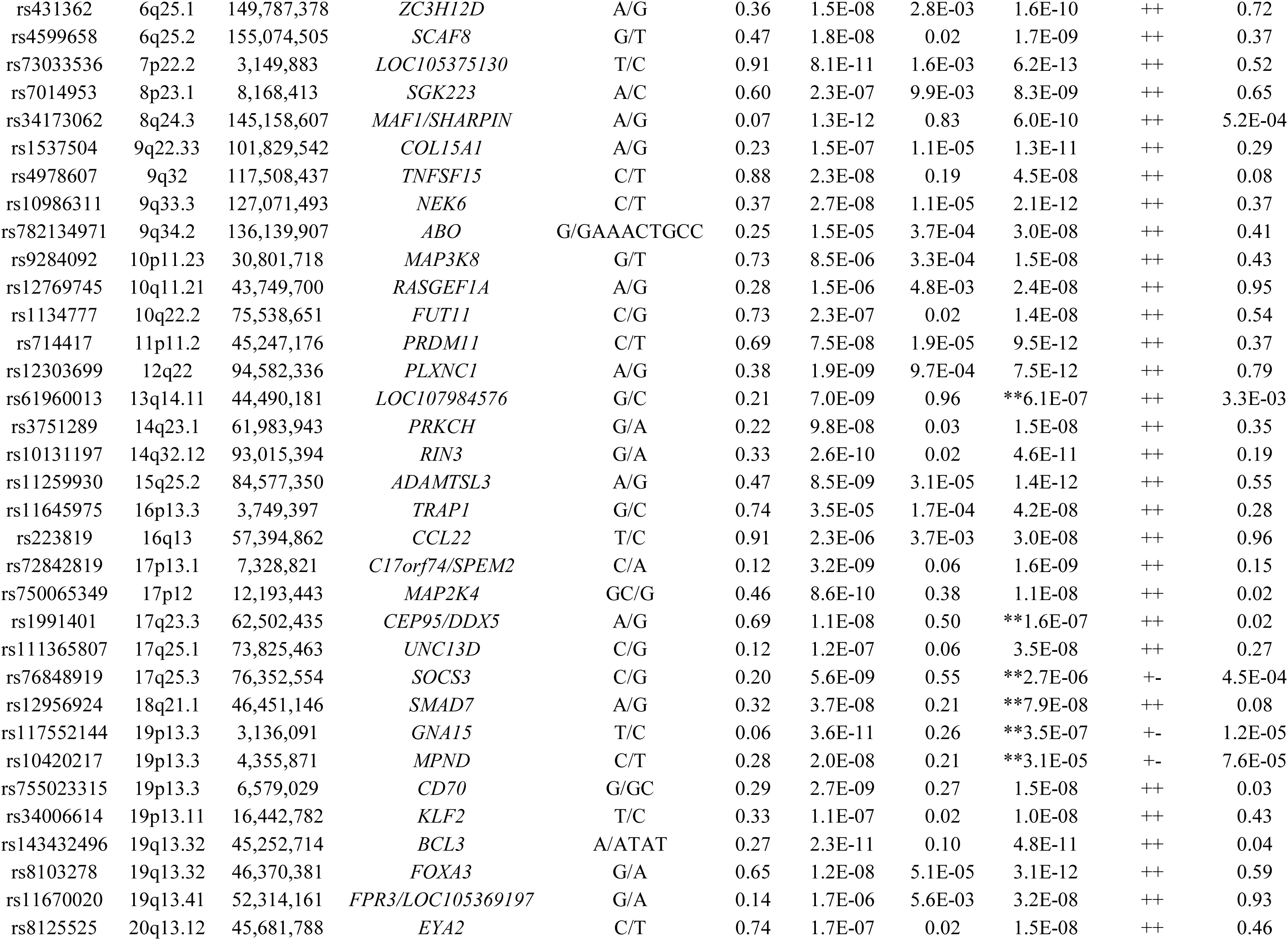
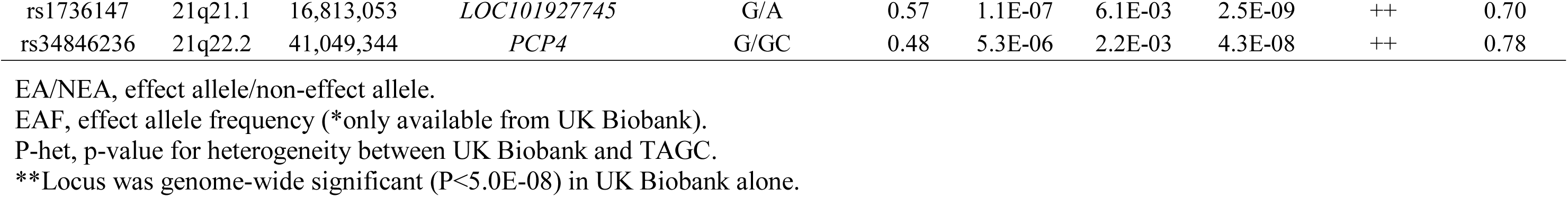
66 novel loci identified for asthma in meta-analysis of UK Biobank and TAGC.

However, despite the P-values not exceeding the genome-wide significance level of 5.0×10^-8^, our meta-analysis did yield evidence for association of 33 of the remaining 36 known asthma loci at a Bonferroni-corrected threshold for testing 145 regions (P=0.05/145=3.4×10^-4^) (**Supplemental Table 4**). The only exceptions were two childhood asthma loci and one for the combined phenotype of asthma and allergic disease (**Supplemental Table 4**). Eight other novel regions were also significantly associated with asthma in UK Biobank alone, but their association signals fell slightly below the genome-wide significance threshold in the meta-analysis (**Fig. 2** and **Table 1**). Thus, our collective analyses with UK Biobank and TAGC identified 66 novel loci for asthma (**Fig. 1**, **Table 1**, and **Supplemental Fig. 3**) and replicated nearly all (142/145) previously known genomic regions. Altogether, the 66 novel loci explained an additional 1.5% of the heritability for asthma and bring the total number of susceptibility loci to 211 at the time of this analysis. Since effect sizes were provided in the publicly available data from TAGC^21^, we also conducted a HapMap-based fixed-effects meta-analysis for asthma with 1,978,494 SNPs common to both datasets and the same number of 536,345 cases and controls. However, no additional novel loci were identified beyond those from the Z-score-based meta-analysis described above.

### Sex-stratified and Polygenic Risk Score Analyses with Asthma Susceptibility Loci

We next used primary level data from UK Biobank to explore how association signals at the 211 asthma susceptibility loci differed in males and females and as a function of cumulative genetic burden. Thirty SNPs yielded nominal P-values for interaction with sex (P-int) that were <0.05, of which only four previously known loci on chromosomes 1p34.3, 2q37.3, 5q22.1, and 6p25.3 would be considered significant at the Bonferroni-corrected threshold for testing 211 loci (P-int=0.05/211=2.4×10^-4^) (**Supplemental Table 5**). Interestingly, the effect sizes at these four loci (and most of the others) were stronger in males than in females, although it should be noted that the significance of the interaction with the chromosome 1p34.3 locus could be inflated due to frequency of the susceptibility allele (**Supplemental Table 5**). We next constructed three weighted genetic risk scores (GRS) in UK Biobank based on the number of risk alleles carried at the 66 novel, 145 known, or all 211 loci. Each GRS was weighted by the overall effect sizes of the included alleles that were derived from the association analysis that included both sexes. As shown in **Fig. 3A**, individuals in the highest decile for the GRS constructed with all 211 risk alleles had nearly a 5-fold increased risk of asthma compared to those in the lowest decile (OR=4.7, 95% CI 4.5-4.9; P<1.0×10^-305^). Increased risk of asthma was also observed for individuals in the highest versus lowest deciles of the GRS for the 145 known and 66 novel loci (**Fig. 3A**), although the overall effect size for the 66-locus GRS was weaker (OR=2.0, 95% CI 1.9-2.1; P=2.6×10^-267^). To determine whether cumulative genetic burden differed in men and women, we also used sex-specific effect sizes to construct weighted GRS in men and women separately for the 66 novel, 145 known, or all 211 asthma loci. This analysis revealed a pattern where risk of asthma among subjects in the highest versus lowest decile of the GRS was increased by approximately one order of magnitude greater in men than in women (Fig. 3B-D). For example, compared to the lowest decile, the OR for asthma among men in the highest decile for the GRS constructed with all 211 loci was 5.5 (95% CI 5.2-5.9; P<1.0×10^-305^) whereas the same analysis in women revealed that the increased risk of asthma among subjects in the highest decile was only 4.4 (95% CI 4.2-4.6; P<1.0×10^-305^). These results were accompanied by significant gene-sex interactions when comparing the highest versus lowest decile for the GRSs constructed from all 211 (P-int=1.2×10^-9^), 145 known (P-int=2.9×10^-9^), and 66 novel (P-int=0.029) loci. Significant interactions with sex were also observed when considering the entire spectrum of cumulative genetic burden for the 211 (P-int=4.0×10^-18^), 145 known (P-int=8.0×10^-18^), and 66 novel (P-int=0.009) loci. The interactions with sex across all deciles remained significant even after exclusion of the four risk alleles on chromosomes 1p34.3, 2q37.3, 5q22.1, and 6p25.3 from the 211-(P-int=2.3×10^-13^) and 145-locus (P-int=1.6×10^-13^) GRSs.

**Figure 3.**
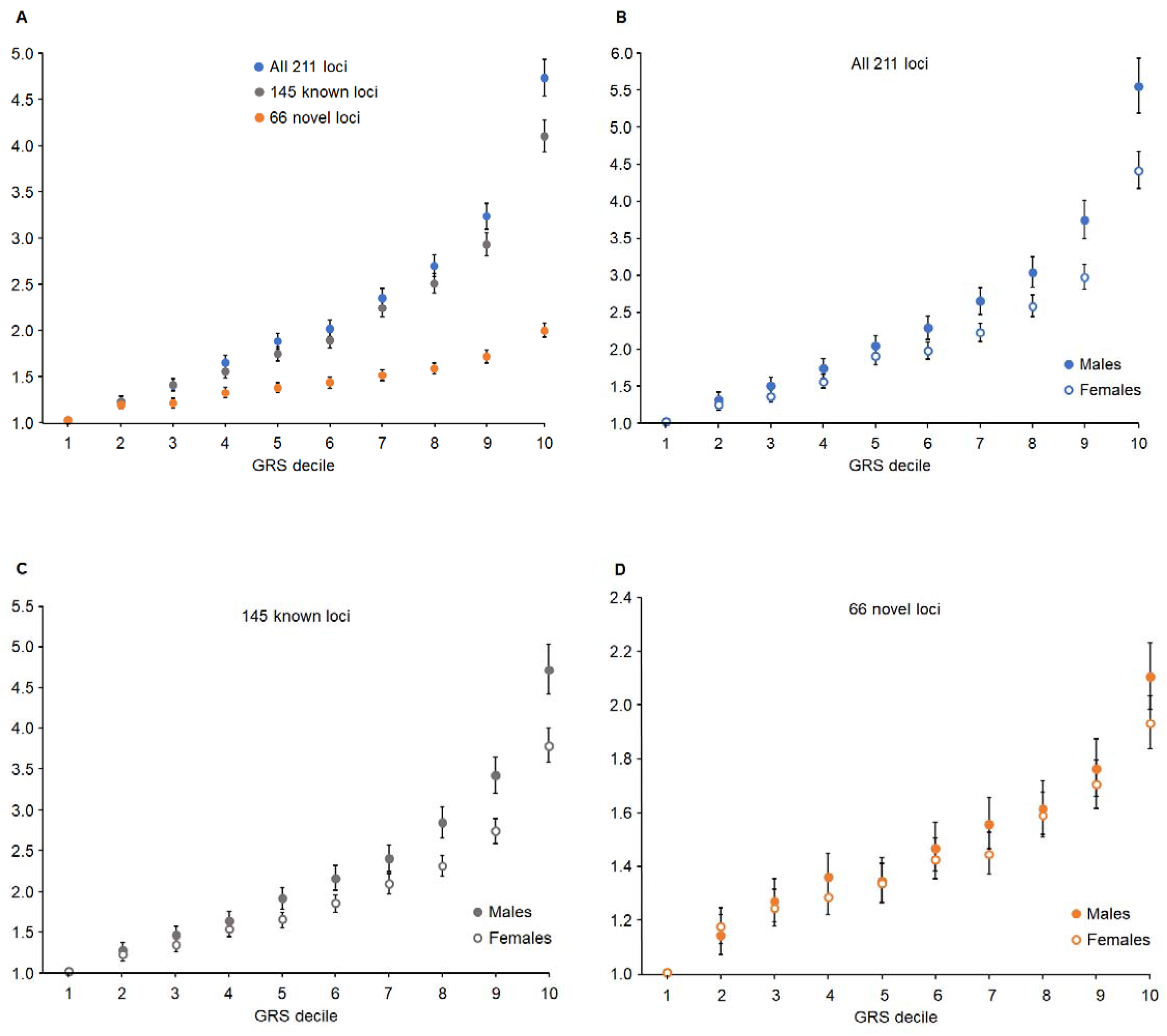
Association of cumulative genetic risk with asthma in UK Biobank. **(A)** Individuals in the highest decile for three weighted polygenic genetic risk scores (GRSs) based on the known or newly identified susceptibility loci have significantly elevated risk for asthma compared the lowest decile. (**B-D**) Compared to females, cumulative genetic risk for asthma was more significantly pronounced in males across the entire spectrum of sex-specific weighted GRSs constructed from all 211 (P-int=4.0×10^-18^), 145 known (P-int=8.0×10^-18^), and 66 novel (P-int=0.009) loci. Interactions with sex were also significant when comparing the highest versus lowest decile for the GRSs constructed from all 211 (P-int=1.2×10^-9^), 145 known (P-int=2.9×10^-9^), and 66 novel (P-int=0.029) loci.

### Association of Novel Loci with Asthma-related Phenotypes and Other Disease Traits

We next used data from UK Biobank to characterize the novel loci with respect to effects sizes for risk of asthma or allergic disease as well as for association with lung function traits. Nearly all 66 novel loci yielded ORs ≤1.10 for association with asthma, with the majority having even more modest effect sizes (**Supplemental Table 6**). Furthermore, most of the novel loci were associated with the more broadly defined phenotype of asthma that also included allergic disease, although some yielded stronger associations with a stricter definition of asthma that excluded allergic disease (**Supplemental Table 6**). We next evaluated the association of all 66 novel asthma loci for associations with forced expiratory volume in one second (FEV_1_) and forced vital capacity (FVC). At the Bonferroni-corrected significance threshold for testing 66 loci (P=0.05/66=7.6×10^-4^), the asthma risk alleles at 17 of the novel loci exhibited directionally consistent associations with decreased FEV_1_, of which 10 were also associated with decreased FVC (**Supplemental Table 6**). To further explore their clinical relevance, we used the PhenoScanner database^31^ to determine whether the novel loci identified in our meta-analyses were associated with other disease phenotypes. Interestingly, lead variants (or tightly linked proxies) at ∼half of the newly identified loci had been previously associated with other asthma-relevant traits, including blood cell parameters (19 loci), height (7 loci), BMI and waist circumference (4 loci), inflammatory bowel disease (3 loci), and cardiovascular disease (1 locus) (**Supplemental Table 7**).

### Enrichment of Asthma-associated Variants in Regulatory Elements or Tissues

To gain biological insight into the newly identified asthma loci, we used two bioinformatics tools to carry out enrichment analyses for regulatory elements and pathways. GWAS Analysis of Regulatory or Functional Information Enrichment with LD correction (GARFIELD) revealed highly significant 3 to 4-fold enrichment of asthma-associated variants colocalizing to DNase I hypersensitive sites in several tissues with biological relevance to asthma, such as, epithelium, lung, and thymus (**Fig. 4A; Supplemental Table 8**). However, enrichment in DNase I hypersensitive elements was particularly evident in blood, and specifically in GM12878 B cells, CD20^+^ B cells, and CD3^+^, CD4^+^, or CD8^+^ T cells (**Fig. 4B; Supplemental Table 8**). A Data-driven Expression-Prioritized Integration for Complex Trait (DEPICT) analysis also yielded consistent results where the most significantly enriched biological pathways and tissues were leukocyte activation/differentiation/proliferation, cytokine production, lymphoid organ development, blood cells, lymphocytes, lymphoid tissue, and spleen (**Supplemental Tables 9 and 10**). Taken together, these results further highlight the immune system as an important component in the pathophysiology of asthma.

**Figure 4.**
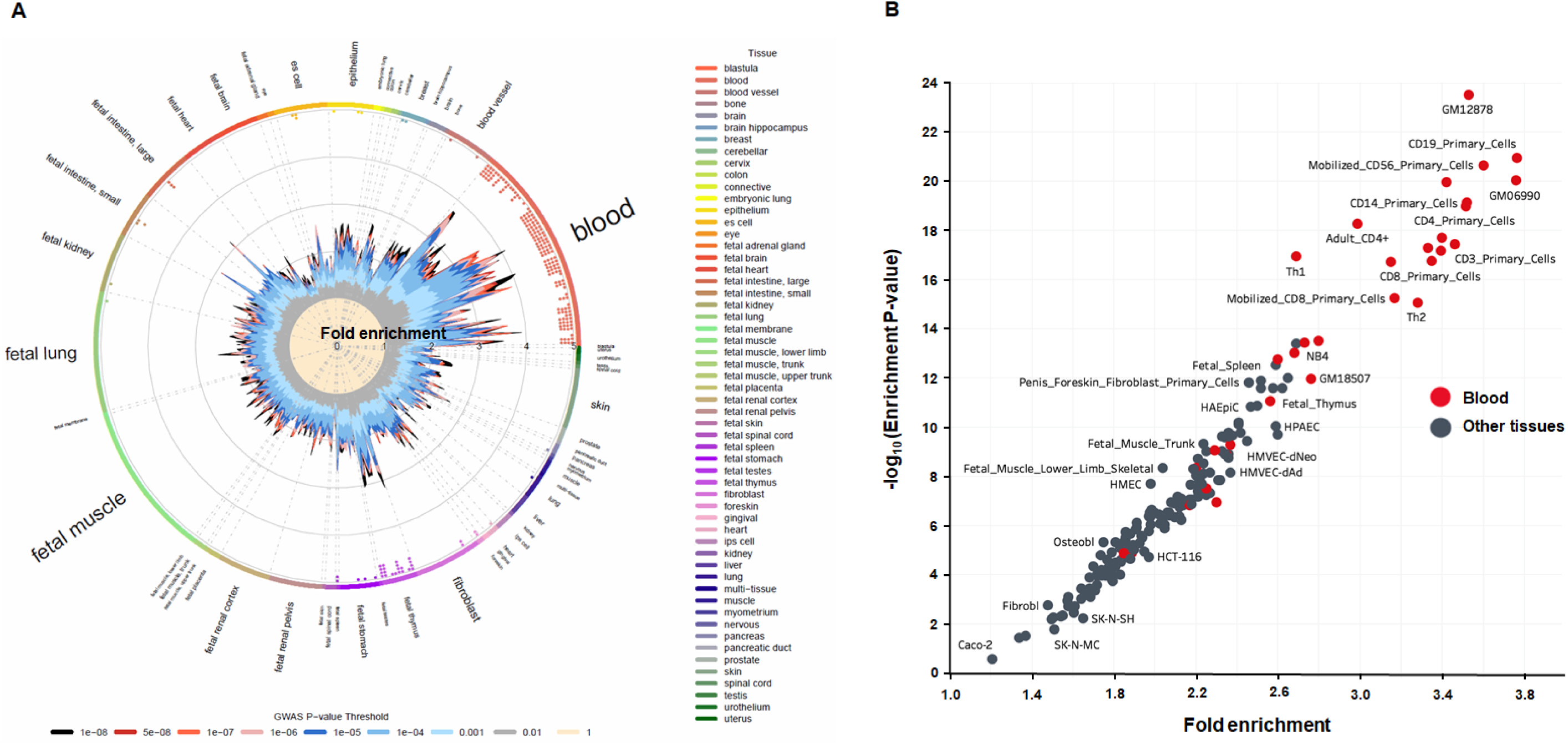
Enrichment of asthma-associated variants in tissue-specific DNase I hypersensitive sites. **(A)** A GARFIELD analysis revealed highly significant 3 to 4-fold enrichment of asthma-associated variants colocalizing to DNase I hypersensitive sites in several asthma-related tissues, such as blood, epithelium, lung, and thymus. The radial plot shows enrichment in the various available tissues using different GWAS significance thresholds for inclusion of asthma-associated variants in the GARFIELD analysis. The small dots on the inside of the outer most circle indicate whether inclusion of asthma-associated variants at GWAS significance thresholds of 1.0×10^−5^ (one dot), 1.0×10^−6^ (two dots), 1.0×10^−7^ (three dots), 1.0×10^−8^ (four dots) in the GARFIELD analysis yielded significant enrichment in that tissue or cell type at P=1.0×10^−15^. **(B)** Highly significant enrichment in DNase I hypersensitive sites was particularly evident in immune cells, such as GM12878 B cells, CD19^+^ B cells as well as CD3^+^, CD4^+^, and CD8^+^ T cells.

### Prioritization of Loci and Positional Candidate Genes

We next used publicly available expression quantitative trait loci (eQTL) data to identify candidate causal genes and prioritize the 66 novel regions for functional validation (**Fig. 1**). The lead SNP or tightly linked (r^2^>0.8) proxy variants at 52 of the novel loci yielded at least one *cis* eQTL for a positional candidate gene in one or more tissues, including those relevant to asthma (**Supplemental Table 11**). Notably, the most significant eQTLs were in blood, some of which yielded P<1.0×10^-300^. These observations are consistent with the GARFIELD enrichment analyses pointing to blood as playing an important role in asthma. We further prioritized loci by focusing on positional candidates that also yielded eQTLs in immune cell populations highlighted by our enrichment analyses, such B and T cells. *CD52*, *AHI1*, and *CLUAP1* on chromosomes 1p36.11, 6q23.3, and 16p13.3 (**Supplemental Fig. 3**), respectively, were three genes that met these criteria and yielded the most significant eQTLs in these specific immune cell types. However, eQTLs were also observed in the lung for *CD52* and *CLUAP1*, making these even stronger candidate causal genes for functional validation (**Supplemental Table 11**). *CLUAP1* was initially identified as a protein that interacts with clusterin and is required for ciliogenesis^32^ but has not otherwise been directly implicated in asthma or pulmonary biology. By comparison, *CD52* encodes a membrane glycoprotein present at high levels on the surface of various immune cells, including lymphocytes, monocytes, and dendritic cells. In addition, alemtuzumab is a monoclonal anti-CD52 (αCD52) antibody that results in preferential and prolonged depletion of circulating T and B cells^33,34^ and is FDA approved for the treatment of relapsing remitting multiple sclerosis^35–37^ and B-cell chronic lymphocytic leukemia^38–40^. Because of its biological role in immune cells relevant to asthma as well as its potential translational implications, we therefore prioritized *CD52* as a strong causal positional candidate gene for functional validation (**Fig. 1**).

### Functional Validation of *CD52*

To functionally validate *CD52*, we used an *in vivo* pharmacological approach. Since alemtuzumab does not target the mouse CD52 protein, we first investigated the immune cell depleting activity of a mouse αCD52 antibody that was previously reported to have biological effects^41^. We designed a mouse experiment that would mimic the clinical protocol for administering alemtuzumab to humans, in conjunction with the induction of airway hyperreactivity through exposure to house dust mite (HDM) (**Supplemental Fig. 4A**). Consistent with the clinical effects of alemtuzumab on circulating lymphocytes in humans^33,34^, the mouse αCD52 antibody efficiently depleted CD4^+^ and CD8^+^ T cells as well as CD19^+^ B cells in both the lung and spleen (**Supplemental Figs. 4B-E** and **5**). We next tested whether αCD52 treatment reduced HDM-induced airway hyperreactivity and lung inflammation (**Fig. 5A**). As expected, mice exposed to HDM had significantly increased lung resistance and decreased dynamic compliance compared to PBS-exposed mice (Fig. 5B and C). However, the induction of lung resistance and deterioration of dynamic compliance as a result of HDM exposure was significantly blunted in mice treated with αCD52 antibody compared to isotype control antibody (Fig. 5B and C). Treatment of HDM-exposed mice with αCD52 antibody also significantly decreased numbers of total cells, eosinophils, T cells, and neutrophils in bronchial alveolar lavage (BAL) (**Fig. 5D** and **Supplemental Fig. 5**). Histological analysis of the lung further showed reduced infiltration of inflammatory cells and decreased thickness of the airway epithelium in HDM-exposed mice treated with αCD52 antibody compared to isotype control antibody (**Fig. 5E**). Collectively, these data demonstrate that targeting CD52 with an antagonizing antibody ameliorates cellular and physiological lung function traits in mice exposed to HDM and provide functional *in vivo* evidence that *CD52* is at least one candidate causal gene at the novel asthma locus on chromosome 1p36.11.

**Figure 5.**
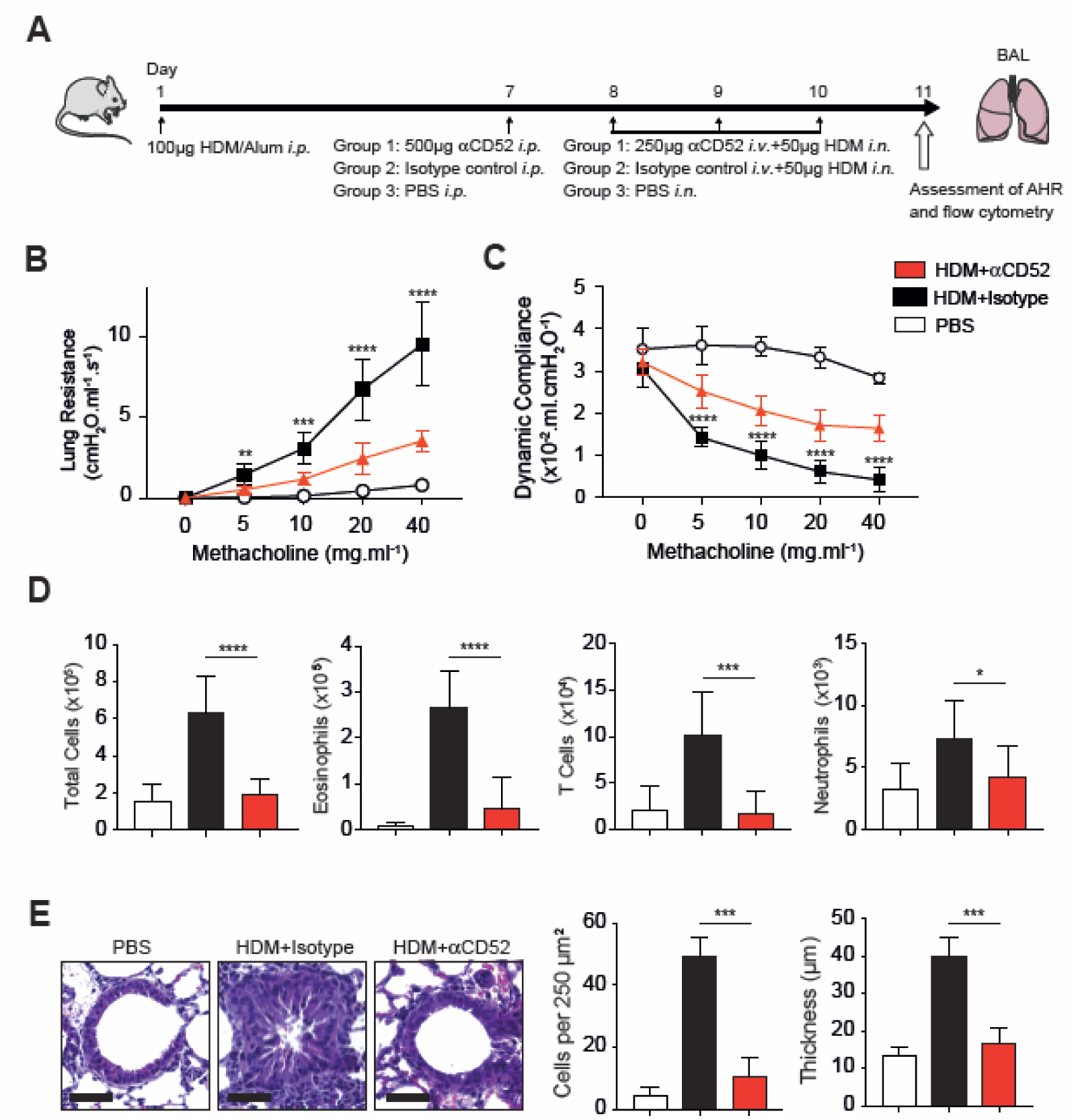
Treatment of mice with anti-CD52 antibody decreases airway hyperreactivity and pulmonary inflammation. (**A)** The experimental protocol for testing the effect of a mouse anti-CD52 (αCD52) antibody on pulmonary function and inflammation was designed to mimic the clinical protocol used to treat multiple sclerosis patients with a human αCD52 antibody (alemtuzumab). Female BALB/cByJ mice were immunized on day 1 with 100µg of house dust mite (HDM) in 2mg of aluminum hydroxide (alum) by intraperitoneal (*i.p.*) injection. On day 7, mice were intraperitoneally administered with either 500µg of αCD52 antibody (Group 1; n=8), 500µg isotype control antibody (Group 2; n=9), or PBS (Group 3; n=4). On days 8, 9 and 10, mice in Groups 1 and 2 were intravenously (*i.v.*) administered 250µg αCD52 antibody or isotype control antibody, respectively, and simultaneously challenged intranasally (*i.n.*) with 50µg HDM. Mice in Group 3 were only challenged intranasally with PBS on days 8, 9 and 10. On day 11, airway hyperreactivity was measured by invasive plethysmography and leukocyte counts were determined by means of flow cytometry. Treatment of mice with αCD52 antibody significantly reduced lung resistance **(B)** and improved dynamic compliance **(C)** in HDM-exposed mice. **(D)** αCD52 antibody significantly decreased numbers of total cells, eosinophils, T cells, and neutrophils in bronchial alveolar lavage (BAL) of mice exposed to HDM compared to the control HDM-exposed isotype antibody group. **(E)** Inflammation and thickness of the airway epithelium was significantly decreased in lungs of HDM-exposed mice treated with αCD52 antibody compared HDM-exposed mice treated with isotype control antibody. Data are shown as mean ± SE. *P<0.05, **P<0.005; ***P<0.0005; ****P<0.0001.

## Discussion

In the present study, we identified 66 novel regions not previously known to be associated with asthma and replicated all but three of the 145 previously identified regions. Despite increasing in the number of genomic regions associated with asthma by nearly 50%, the newly identified 66 loci only explained an additional 1.5% of the heritability for asthma, reflecting the modest nature of the susceptibility alleles’ effect sizes on disease risk. Taking into consideration the ∼7% attributable to previously known loci, the results of our genetic analyses and the 211 susceptibility loci identified to date still only explain 8-9% of the total heritability for asthma. Compared to other common diseases^42^, the relatively low fraction of the heritability explained by genetic factors may be due to the more complex underlying phenotypic and genetic heterogeneity of asthma. For example, most of the novel asthma loci also exhibited associations with allergic disease, consistent with recent studies demonstrating that these phenotypes share genetic determinants^20^. By comparison, only ∼25% of the novel regions exhibited associations with FEV_1_ and FVC. This is the proportion of asthma susceptibility loci that studies have consistently shown to be associated with FEV_1_ and FVC and suggest that most genetic factors predisposing to asthma do not necessarily manifest through associations with lung function traits.

GxE interactions are also recognized as important components of asthma’s complex genetic etiology but progress in this area been limited, with most studies focusing on exposures such as allergens, cigarette smoke, and air pollutants^43,44^. Our data suggest that sex is another potentially important endogenous environmental factor that can modulate risk through interactions with asthma susceptibility alleles, both individually or as a function of cumulative genetic burden. This was particularly evident among men in whom increased risk of asthma across all categorized risk alleles was consistently more accentuated than in females, even after exclusion of four susceptibility alleles that individually yielded statistical evidence for being more strongly associated with asthma in men. Evidence for such gene-sex interactions has been demonstrated for several complex phenotypes^45^, including anthropometric traits^46–48^, coronary artery disease^49^, as well as asthma^50–53^. However, the previously reported sex-specific associations for respiratory outcomes^50–53^, which were identified in much smaller sample sizes relative to our study, were not detected in our present analyses (data not shown). Interestingly, the male-specific locus (rs2549003) identified on chromosome 5^52^ was associated with asthma in our meta-analysis (overall P=2.0×10^-14^) but the association signal was derived equally in men and women. Nonetheless, these data collectively suggest that GxE interactions are likely to have important contributions to risk of asthma and represent an unexplored area for future investigation, particularly as new statistical methods are developed to carry out such analyses on a genome-wide level^54^.

Our results also add to the growing body of genetic and bioinformatics evidence that immune system-related processes are important drivers of asthma pathophysiology, consistent with prior studies^21^. For example, enrichment analyses highlighted pathways related to various aspects of leukocyte function and immune system regulation/development. Furthermore, asthma-associated variants were highly enriched in regions of open chromatin in several tissues and especially in B and T lymphocytes. This concept is also supported by the nature of specific candidate causal genes located at some of the novel loci. One such example is the chromosome 1p36.22 locus where the lead SNP (rs2230624; G>A) yielded the strongest effect size for asthma (OR=1.19) out of all 211 risk variants. Rs2230624 leads to a rare (∼1% frequency in European populations) but computationally predicted deleterious Cys273Tyr substitution in exon 8 of *TNFRSF8*, which encodes the costimulatory molecule CD30. However, the common Cys273 allele of rs2230624 is associated with increased risk, suggesting that loss of CD30 function may be protective for asthma. This hypothesis would be consistent with prior studies showing that CD30 increases pro-inflammatory cytokine production by CD4^+^ T cells and is involved in Th1, Th2, and Th17 immune responses^55^. Furthermore, soluble CD30 levels are elevated in patients with asthma or atopic dermatitis and correlate with disease severity^56,57^. When gene expression data were used to prioritize candidate causal genes, *CD200R1* and *AHI1* emerged as two other potential genes of interest related to immune function, given the highly significant *cis* eQTLs (P<1.0×10^-300^) observed in blood with the lead variants on chromosomes 3q13.2 (rs7622814) and 6q23.3 (rs4526212), respectively. For example, CD200:CD200R1 signaling suppresses expression of proinflammatory molecules^58^ and the induction of CD8^+^ T cells^59^. By comparison, *AHI1* may be involved in JAK-STAT signaling^60^ and downstreatm effects on Th1 and Th2 cells^61^. In addition, a recent GWAS identified a variant (rs9647635) at the chromosome 6q23.3 locus in near perfect LD with our peak SNP (r^2^=0.98) that was associated with selective IgA deficiency^62^, which is correlated with asthma^63^.

Another important aspect of our study are the *in vivo* pharmacological experiments functionally validating *CD52* as at least one candidate causal gene at the chromosome 1p36.11 locus. These findings not only point to a novel pathway for development of asthma but may also have potential translational implications. For example, we demonstrated that treatment of mice with an αCD52 antibody mimicked the immune cell-depleting activity of alemtuzumab in the circulation in humans^33,34^. We readily observed depletion of immune cells in spleen, which in mice is considered as accurately reflecting the pool of peripheral immune cells^64^. By also depleted leukocytes in lung and BAL, the mouse αCD52 antibody had biological effects in tissues that, to our knowledge, have not been reported with alemtuzumab in humans. Of most importance and relevance to asthma, mice receiving the αCD52 antibody exhibited significantly reduced pulmonary inflammation, airway epithelium thickness, and allergen-induced airway hyperreactivity. Given that alemtuzumab is already FDA-approved^35–40^, it is tempting to consider CD52 as a novel therapeutic target for asthma. However, serious, albeit very rare, side effects have been reported with alemtuzumab, and as an immunosuppressive agent, patients being treated with this monoclonal antibody are at increased risk of infections and malignancies^65^. Nonetheless, there may still be a population of severe asthmatics with limited therapeutic options^66^, such as those with corticosteroid-resistant or eosinophilic asthma, in whom the clinical benefits of alemtuzumab may justify its potential adverse effects. The generally favorable long-term outcomes observed with alemtuzumab in small numbers of patients with hypereosinophilic syndrome^67,68^ suggest this treatment strategy may be a potentially viable concept worthy of further consideration. Approaches to limiting or eliminating serious adverse effects of alemtuzumab could be through reduced dosing strategies, such as those used for multiple sclerosis compared to leukemia^69^, and development of other αCD52 antibodies that may be more efficacious but less immunogenic^70^. Alternatively, the precise mechanisms through which alemtuzumab exerts its therapeutic effects are still not entirely known. Thus, a better understanding of CD52’s biological functions may identify alternative targets that could be evaluated for therapeutic development and which may lead to less severe or minimal side effects.

While the present analyses reveal novel genetic determinants of asthma, our study should also be taken in the context of certain limitations. First, our inclusion of subjects in UK Biobank with self-reported asthma could have led to misclassification of cases and biased the results towards the null. However, the large sample size in our study and our ability to replicate all but three of the known 145 asthma loci indicate that our analyses were robust to the presence of such potential confounding factors. Second, most subjects in UK Biobank and TAGC were of European ancestry and it is possible that the genetic association results may not be generalizable to other populations. This notion has previously been observed with both common and rare variants even among closely related Latino populations and in comparison, to subjects of African ancestry^27,71^. Third, the asthma loci identified in our meta-analysis relied on GWAS results in TAGC that were derived from imputed Z scores and P-values. Although this approach allowed us to include the largest number of SNPs possible in the meta-analysis with UK Biobank, it may not have provided the most precise summary statistics in TAGC. The likelihood of this potential limitation affecting our overall results is low since the Z-score meta-analysis replicated the association signals at nearly all known asthma susceptibility loci. Moreover, even though the Z-score meta-analysis did not provide effect sizes, the ORs based on association tests in UK Biobank alone are still likely to be very similar to those derived from the combined datasets given the large size of UK Biobank. Lastly, our study was primarily focused on discovery of main genetic effects of common susceptibility alleles. However, rare variants or GxE interactions, particularly with respect to other environmental exposures besides sex, may still play important roles in modulating asthma risk and will need to be addressed in appropriately designed future studies.

In summary, large-scale genetics analyses identified 66 novel risk loci for asthma and implicated regulation of immune processes as an important component to disease susceptibility. These results provide opportunities for further exploration of the biological mechanisms underlying the pathogenesis of asthma and evaluating additional validated causal genes as potentially novel therapeutic targets.

## Materials and Methods

### Study Populations

UK Biobank recruited participants between 40-69 years of age who were registered with a general practitioner of the UK National Health Service (NHS). Between 2006-2010, a total 503,325 individuals were included. All study participants provided informed consent and the study was approved by the North West Multi-centre Research Ethics Committee. Detailed methods used by UK Biobank have been described elsewhere^72^. Participating cohorts in the Trans-National Asthma Genetic Consortium (TAGC) have been previously described in detail^30^. Briefly, TAGC includes 56 studies of European populations, 7 studies of African populations, 2 studies of Japanese populations and one study of a Latino population. All participants gave written consent for participation in genetic studies, and the protocol of each study was approved by the corresponding local research ethics committee or institutional review board. The present study was approved by the Institutional Review Boards of the USC Keck School of Medicine.

### GWAS Analyses in UK Biobank

Asthma cases was defined based on UK Biobank field code 6152_8 (doctor diagnosed asthma), International Classification of Diseases version-10 (ICD10) J45 (asthma)/J46 (severe asthma), and self-reported asthma. Field 6152 is a summary of the distinct main diagnosis codes a participant has had recorded across all their hospital visits. Non-asthmatic controls were defined as individuals free from 6152_8 (doctor diagnosed asthma), 6152_9 (doctor diagnosed allergic disease), ICD10 J45/J46/J30 (hay fever)/L20 (dermatitis and eczema), as well as self-reported asthma/hay fever/eczema/allergy/allergy to house dust mite (HDM). This definition strategy resulted in 64,538 asthma cases and 329,321 controls in UK Biobank, and was selected to increase the likelihood of identifying loci associated specifically with asthma rather than loci for asthma and/or allergic disease. Quality control of samples and DNA variants, and imputation were performed by the Wellcome Trust Centre for Human Genetics, as described in detail elsewhere^72^. Briefly, ∼90 million SNPs imputed from the Haplotype Reference Consortium, UK10K, and 1000 Genomes imputation were available in UK Biobank. Of these, 9,572,556 variants were used for GWAS analysis after filtering on autosomal SNPs with INFO scores >0.8 (directly from UK Biobank) and with minor allele frequencies (MAF) >1% in the 487,409 individuals with imputed genotypes. A GWAS analysis was performed with BOLT-LMM v2.3.2 using a standard (infinitesimal) mixed model to correct for structure due to relatedness, ancestral heterogeneity, with adjustment for age, sex, the first 20 principal components, and genotyping array^73^. The genome-wide significance threshold was set at P=5.0×10^-8^. Since BOLT-LMM relies on linear models even for qualitative traits, SNP effect size estimates on the quantitative scale were transformed to obtain odds ratios (ORs) and standard errors (SEs) using the following formula: β or SE/(μ ∗ (1 − μ)), where μ = case fraction^73^.

### Imputation in TAGC

Publicly available GWAS summary statistics for asthma with 2,001,281 SNPs in predominately European ancestry populations from TAGC were downloaded and imputed with summary statistics imputation (SSimp) software^74^ using the European population from the 1000 Genomes Project (Phase 1 release, v3) as a reference panel for LD computation. This imputation resulted in Z-scores and P-values for association of 9,415,011 SNPs with asthma in TAGC.

### Meta-analyses for asthma in UK Biobank and TAGC

The imputed Z-score and P-value summary level data in TAGC were combined with the results of our linear mixed model GWAS for asthma in UK Biobank. We first performed a Z-score meta-analysis for asthma with 8,365,359 SNPs common to both datasets assuming an additive model, as implemented in METAL^75^. We also performed a fixed effect meta-analysis for asthma with betas and standard errors obtained in our GWAS analysis of UK Biobank and those provided in TAGC for 1,978,494 SNPs common to both datasets. The genome-wide threshold for significant association in both meta-analyses was set at P=5.0×10^-8^. A locus was defined as novel if our sentinel SNP was >1Mb away or in weak or no linkage disequilibrium (r^2^≤0.1) with the lead variants at the 145 previously reported loci for asthma and/or allergic disease^5–11,13,15–21,23,24^. Replication of the known asthma/ allergic disease loci in our meta-analysis was considered significant at a Bonferroni-corrected threshold of P=3.4×10^-4^ for testing 145 loci (0.05/145).

### Sex-stratified associations with asthma susceptibility loci

Sex-stratified association with variants at all 211 loci and risk of asthma was carried out with primary level data in UK Biobank using logistic regression in males and females separately, adjusted for age, the first 20 principal components, and genotyping array. Logistic regression with the inclusion of a SNP * sex interaction term was also used to test for an interaction between SNP and sex on risk of asthma according to the following model: Logit [p] = β_0_ + β_1_ SEX + β_2_ SNP + β_3_ SEX * SNP. Interaction P-values (P-int) were considered significant at the Bonferroni-corrected threshold for testing 211 loci (0.05/211=2.4×10^-4^). Sex-stratified and gene-sex interaction analyses were performed with STATA (v15.0, StataCorp LP, Texas, USA).

### Evaluation of asthma loci as a function of cumulative genetic burden

Primary level data from UK Biobank were also used to generate weighted genetic risk scores (GRSs) for the 66 novel, 145 known, or all 211 loci. For each variant, the number of risk alleles was multiplied by its respective weight (i.e., the Ln of the OR) obtained from the GWAS analysis carried out in UK Biobank and summed together across all variants to generate the three GRSs. Sex-specific weighted GRSs were also constructed for the 66 novel, 145 known, or all 211 loci using effect sizes obtained from the logistic regressions carried out in males and female separately. Association between weighted GRSs and asthma in all subjects and in males and females separately was tested by classifying participants into deciles according to the distribution of weighted GRS and using logistic regression to calculate ORs for subjects in the highest decile compared to the lowest decile, with adjustment for age, sex (if applicable), the first 20 principal components, and genotyping array. Tests for an interaction between sex and the GRS on risk of asthma were carried out by including a sex * sex-specific GRS interaction term in the logistic regression comparing the highest versus lowest decile or by including the sex-interaction term in a model across all deciles, with inclusion of the same covariates. Sex-stratified and gene-sex interaction analyses were performed with STATA (v15.0, StataCorp LP, Texas, USA).

### Association of novel loci with asthma-specific phenotypes, allergic-specific phenotypes, lung function, and other disease traits

Association signals at the novel loci were characterized further by testing the lead SNPs for association with asthma-vs. allergic disease-specific phenotypes as well as lung function traits in UK Biobank. Asthma-specific cases were defined as positive for 6152_8 (doctor diagnosed asthma) or ICD10 J45 (asthma)/J46 (severe asthma) or self-reported asthma and negative for 6152_9 (doctor diagnosed allergic disease), ICD10 J30 (hay fever) /L20 (dermatitis and eczema), and self-reported hay fever/eczema/allergy/allergy to house dust mite. Allergic disease-specific cases were defined as positive for 6152_9 (doctor diagnosed allergic disease), ICD10 J30 (hay fever) /L20 (dermatitis and eczema), or self-reported hay fever/eczema/allergy/allergy to house dust mite and negative for 6152_8 (doctor diagnosed asthma), ICD10 J45/J46, and self-reported asthma. Controls were defined as individuals free from 6152_8 (doctor diagnosed asthma), 6152_9 (doctor diagnosed allergic disease), ICD10 J45/J46/J30/L20, and self-reported hay fever/eczema/allergy/allergy to HDM. These definitions lead to the inclusion of 33,830 cases/329,321 controls for the asthma-specific phenotype and 93,468 cases/329,321 controls for the allergic disease-specific phenotype in UK Biobank. The lead SNPs were also tested for association with forced expiratory volume in 1-second spirometry (FEV_1_; field codes 3063, N=446,697) and forced vital capacity spirometry (FVC, field code 3062, N=446,697). Both lung function traits were log-transformed prior to analysis and a Bonferroni-corrected P=7.6×10^-4^ (0.05/66) for testing 66 loci was considered as the threshold for significant association with FEV_1_ and FVC. Analyses were performed with BOLT-LMM V2.3.2 using a standard (infinitesimal) mixed model to correct structure due to relatedness, ancestral heterogeneity, or other factors, with adjustment for age, sex, the first 20 principal components, and genotyping array^73^. Evaluation of the novel asthma loci for association with other disease traits was carried out using the Phenoscanner database^31^. Only associations with P<5.0×10^-8^ and derived from SNPs in high LD (r^2^>0.8) with our lead GWAS variants are reported.

### Enrichment of asthma loci in epigenetic marks

Enrichment of asthma-associated variants in DNase I hypersensitive sites (peaks) was determined using the GWAS Analysis of Regulatory or Functional Information Enrichment with LD correction (GARFIELD) method^76^. Briefly, GARFIELD leverages GWAS findings with regulatory or functional annotations (primarily from ENCODE and Roadmap epigenomics data)^77^ to find features relevant to a phenotype of interest. It performs greedy pruning of GWAS SNPs (r^2^>0.1) and then annotates them based on functional information overlap. Next, it quantifies ORs at various significance cutoffs and assesses them by employing a generalized linear model framework, while matching for minor allele frequency, distance to nearest transcription start site and number of LD proxies (r^2^>0.8).

### Data-driven Expression-Prioritized Integration for Complex Traits (DEPICT)

Additional gene prioritization analysis and tissue enrichment analysis were carried out by DEPICT (version 1.1)^78^ using 33,017 genome-wide significant SNPs (P<5.0×10^-8^) associated with asthma that were identified in the meta-analysis with UK Biobank and TAGC. Both nominal P-values and false discovery rates (FDRs) were calculated for gene set enrichment and tissue enrichment.

### Heritability

Heritability was calculated with the INDI-V calculator under multifactorial liability threshold model. The baseline population risk (K) was set at 15%^25^ and twin heritability (h_L_^2^) was set at 65% (average of 35% and 95%)^3^. GWAS results from UK Biobank were used to estimate the heritability attributable to asthma-associated variants at the 66 novel loci.

### Expression Quantitative Trait Locus (eQTL) analyses

Functional evaluation of SNPs at the 66 novel asthma loci and prioritization of candidate causal genes was determined using multi-tissue eQTL data from the GTEx Project (version 6)^79^ and the PhenoScanner database^31^. Consideration was only given to *cis* eQTLs with P<5.0×10^−8^ that were derived from our lead GWAS variants or proxy SNPs in high LD (r^2^>0.8).

### *In vivo* evaluation of airway hyperreactivity after pharmacological perturbation of CD52

Female BALB/cByJ mice between 6-8 weeks of age were immunized on day 1 with 100µg of house dust mite (HDM) in 2mg of aluminum hydroxide (alum) by intraperitoneal (*i.p.*) injection. On day 7, mice were randomly assigned to three groups. Group 1 was intraperitoneally administered 500µg of monoclonal αCD52 IgG2a antibody (clone BTG-2G, MBL International, Woburn, MA) and Group 2 was intraperitoneally administered 500µg IgG2a isotype control antibody (clone 2A3, BioXCell, West Lebanon, NH). Group 3 only received PBS as a control. On days 8, 9 and 10, mice in Groups 1 and 2 were intravenously (*i.v.*) administered 250µg of the αCD52 antibody or isotype control antibody, respectively, and simultaneously challenged intranasally (*i.n.*) with 50µg HDM. Mice in Group 3 were only challenged intranasally with PBS on days 8, 9 and 10. On day 11, airway hyperreactivity was measured by invasive plethysmography. Lung resistance and dynamic compliance at baseline and in response to sequentially increasing doses of methacholine (5-40mg/ml) was measured using a Buxco FinePointe (Data Sciences International, St. Paul, MN) respiratory system in tracheostomized immobilized mice that were mechanically ventilated under general anesthesia, as described elsewhere^80^. After airway hyperreactivity measurements were completed, the trachea was cannulated and 2ml of PBS was used to collected bronchial alveolar lavage (BAL) fluid for analysis of immune cells by flow cytometry. All animal studies were approved by the USC Keck School of Medicine Institutional Animal Care and Use Committee and conducted in accordance with the Department of Animal Resources’ guidelines.

### Flow Cytometry

Single cell suspensions were prepared from lung and spleen tissue in accordance with standard protocols. Immune cells were quantified by means of flow cytometry after staining cells with phycoerythrin (PE)-anti-CD52 (clone BTG-2G, MBL International, Woburn, MA), allophycocyanin (APC)/Cy7-anti-CD45 (clone 30-F11), fluorescein isothiocyanate (FITC)-anti-CD19 (clone MB19-1), peridinin-chlorophyll-protein complex (PerCP)/Cy5.5-anti-CD3ε (clone 17A2), brilliant violet 421™ (BV)-anti-CD4 (clone GK1.5), PE/Cy7-anti-CD8 (clone 53-6.7), (BioLegend, San Diego, CA) in the presence of anti-mouse FC-block (clone 2.4G2, BioXcell, West Lebanon, NH). Leukocytes in BAL fluid were quantified after staining cells with PE-anti-Siglec-F (clone E50-2440, BD Biosciences, San Jose, CA), PE/Cy7-anti-CD45 (clone 30-F11), APC/Cy7-anti-CD11c (clone N418), FITC-anti-CD19 (clone MB19-1), PerCP/Cy5.5-anti-CD3ε (clone 17A2), APC-anti-Gr-1 (clone RB6-8C5) (BioLegend, San Diego, CA), and eFluor450-anti-CD11b (clone M1/70, eBioscience, San Diego, CA) in the presence of anti-mouse FC-block (clone 2.4G2, BioXcell, West Lebanon, NH). The gating strategy used with these antibodies is shown in **Supplemental Fig. 5**. CountBright absolute count beads (Thermo Fisher Scientific, Waltham, Mass) were used to calculate absolute cell number, according to the manufacturer’s instructions. At least 10^5^ CD45^+^+ cells were acquired on a BD FACSCanto II (BD Biosciences). Data were analyzed with FlowJo software (TreeStar, Ashland, OR).

### Histology

Lungs were harvested, fixed overnight in 4% paraformaldehyde, and embedded in paraffin. 4µm sections were cut and stained with H&E according to standard protocols. Images of the H&E-stained slides were acquired with a KeyenceBZ-9000 microscope (Keyence, Itasca, Ill) and analyzed for the number of inflammatory cells and thickness of the airway epithelium with the ImageJ Analysis Application (NIH & LOCI, University of Wisconsin).

### Statistical Analysis

Differences in measured variables between groups of mice were determined by unpaired Student’s t-test (SAS 9.3; SAS Institute Inc). Data are expressed as mean ± SE and differences were considered statistically significant at P<0.05.

### URLs

UK Biobank, http://www.ukbiobank.ac.uk/; Trans-National Asthma Genetic Consortium, https://www.ebi.ac.uk/gwas/downloads/summary-statistics; Genotype-Tissue Expression Portal, http://gtexportal.org/; Phenoscanner, http://www.phenoscanner.medschl.cam.ac.uk/phenoscanner; GARFIELD, https://www.ebi.ac.uk/birney-srv/GARFIELD/; DEPICT, http://www.broadinstitute.org/mpg/depict; R statistical software, http://www.R-project.org/;

## Data Availability

Full summary statistics relating to the GWAS meta-analysis will be deposited with The NHGRI-EBI Catalog of published genome-wide association studies: https://www.ebi.ac.uk/gwas/. All other relevant data will be made available by the corresponding author upon reasonable request.

## Acknowledgments

This work was supported, in part, by National Institutes of Health Grants R01HL133169, R01ES021801, R01ES025786, R01ES022282, R21ES024707, R21AI109059, P30ES007048, and P01ES022845, and U.S. EPA Grant RD83544101. The funders had no role in study design, data collection and analysis, decision to publish, or preparation of the manuscript.

## Author Contributions

Concept and design: Y.H., J.A.H., and H.A. Acquisition, analysis, and interpretation of data: Y.H., Q.J., P.S.J., B.P.H., C.P., P.H., J.G., N.C.W., E.E., F.G., O.A., J.AH., and H.A. Drafting of the manuscript: Y.H., J.H., and H.A. Critical revision of the manuscript for important intellectual content: All authors.

