## Supplemental Materials for "Large-scale Genetic Analysis Identifies 66 Novel Loci for Asthma"

Han et al.

### Address correspondence and reprint requests to:

Hooman Allayee, PhD

Department of Preventive Medicine Keck School of Medicine of USC 2250 Alcazar Street, CSC202

Ema

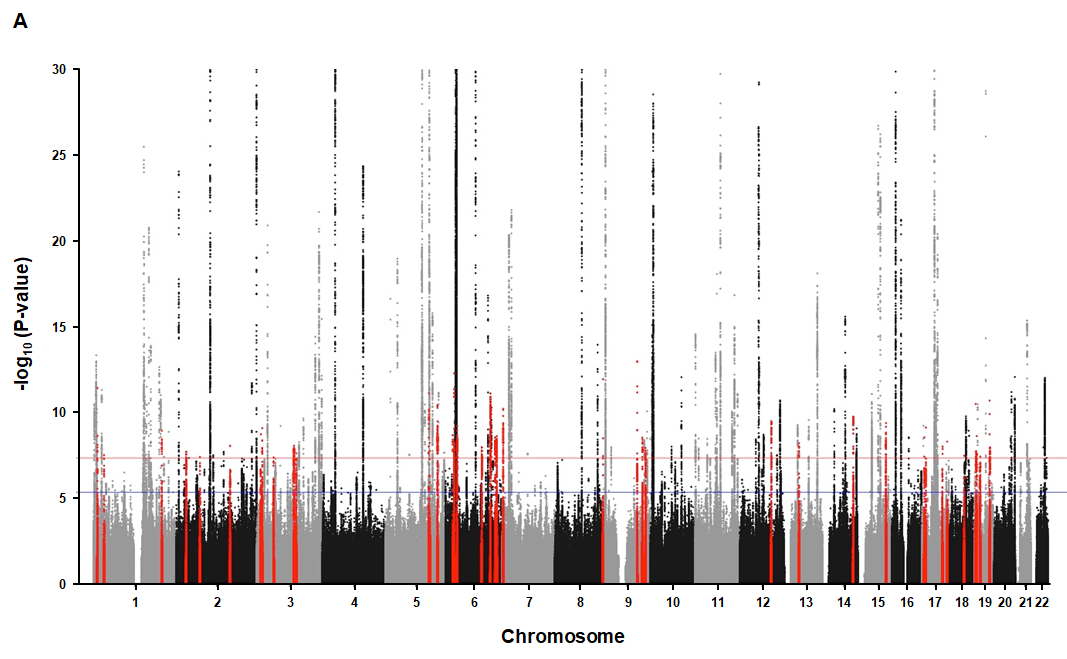

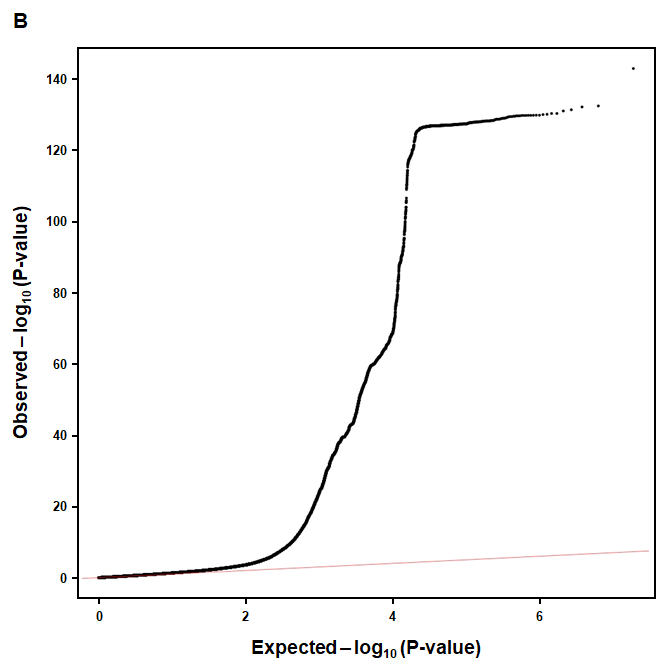

**Supplemental Figure 1. Results of GWAS analysis for asthma in UK Biobank. (A)** A Manhattan plot shows 41 novel loci (red dots) significantly associated with asthma in UK Biobank. The GWAS analysis included 64,538 cases and 329,321 controls. Genome-wide thresholds for significant (P=5.0x10^-8^) and suggestive (P=5.0x10^-6^) association are indicated by the horizontal red and dark blue lines, respectively. P-values are truncated at -log_10_(P)=30. **(B)** A quantile-quantile plot shows the observed versus the expected P-values from the association analyses for asthma in UK Biobank. Although the genomic control factor (λ) in UK Biobank was 1.31, the LD Score intercept from BOLT-LMM was 1.076 (SE=0.009), suggesting there was no inflation due to population structure.

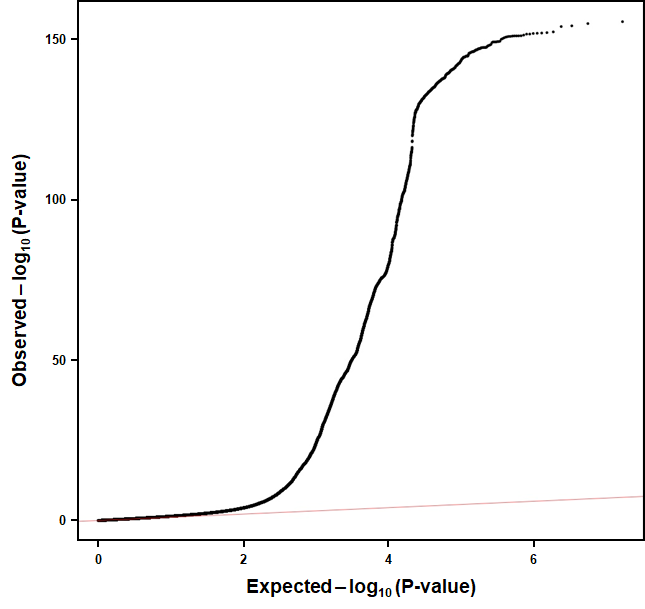

**Supplementary Figure 2. Quantile-quantile plots for results of GWAS meta-analysis with UK Biobank and TAGC.** The observed versus the expected P-values from the Z-score meta-analysis are shown. The meta-analysis for asthma included a total of 88,486 cases and 447,859 controls from UK Biobank (64,538 asthma cases and 329,321 controls) and TAGC (23,948 asthma cases and 118,538 controls) and 8,365,715 SNPs common to both datasets.

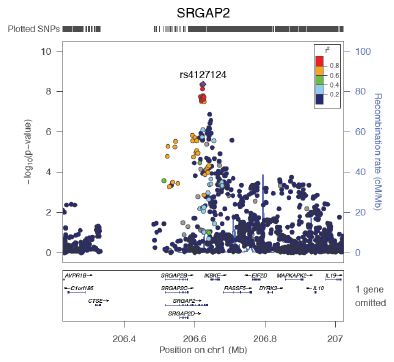

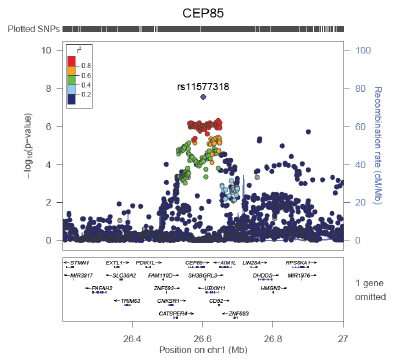

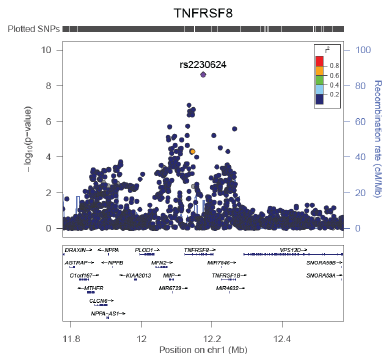

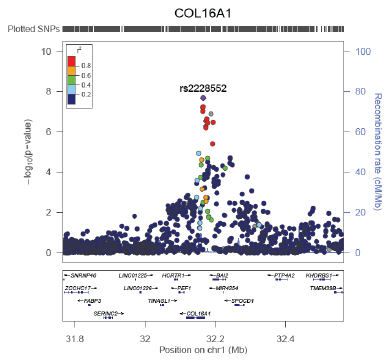

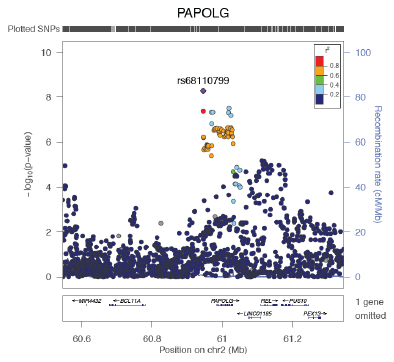

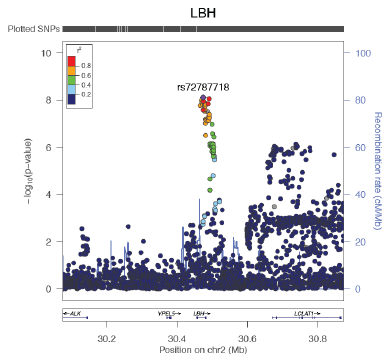

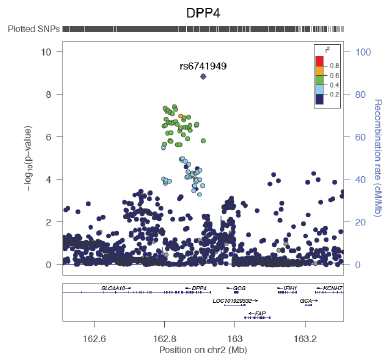

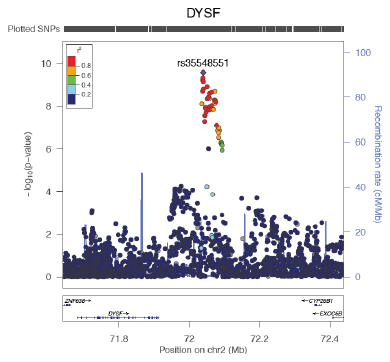

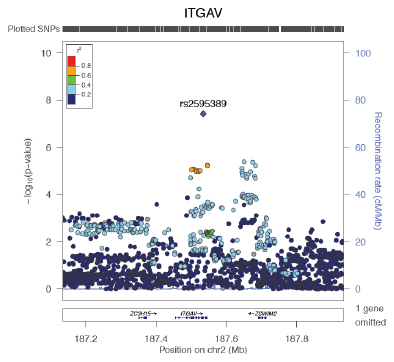

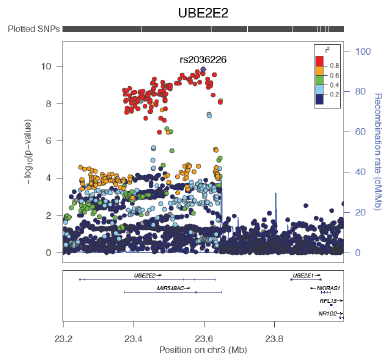

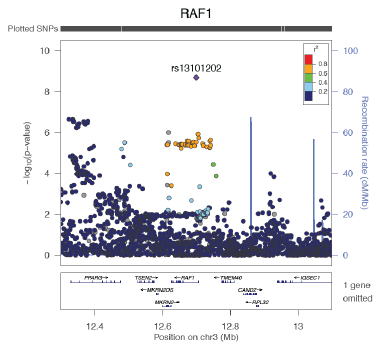

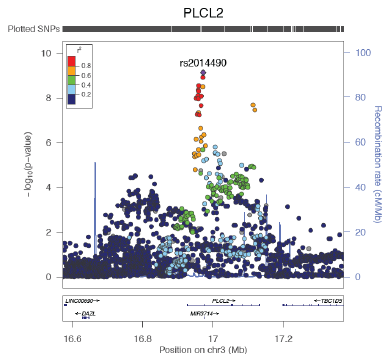

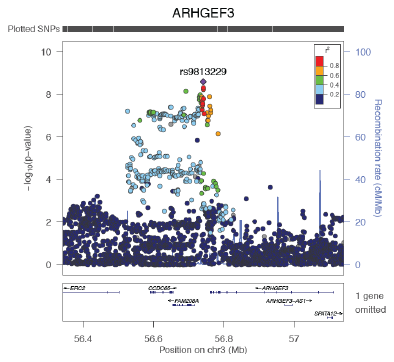

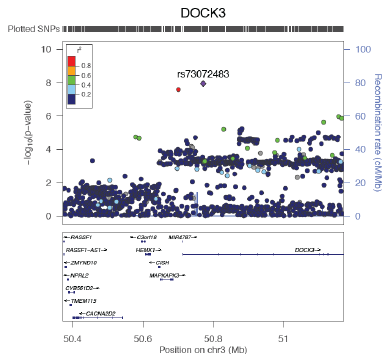

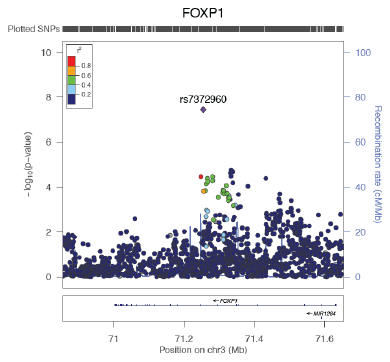

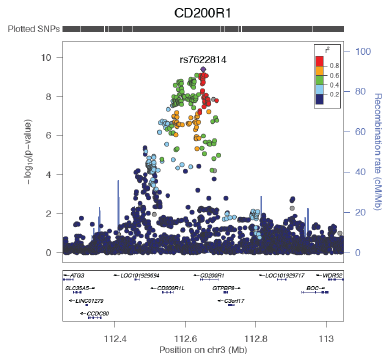

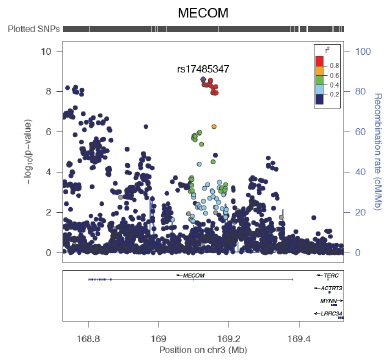

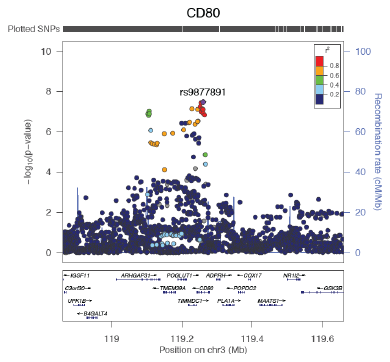

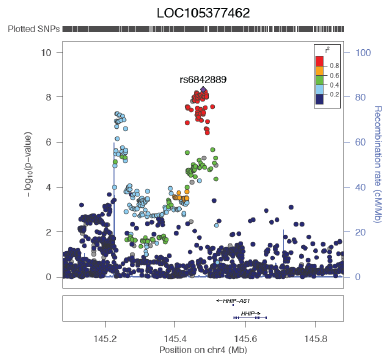

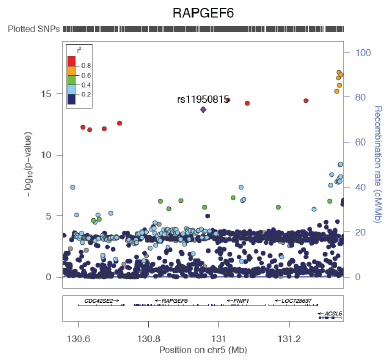

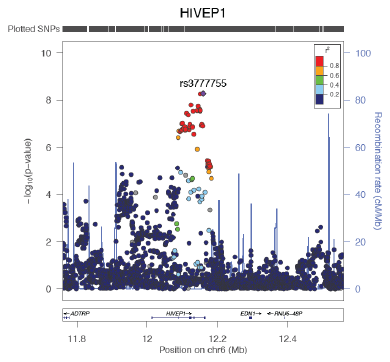

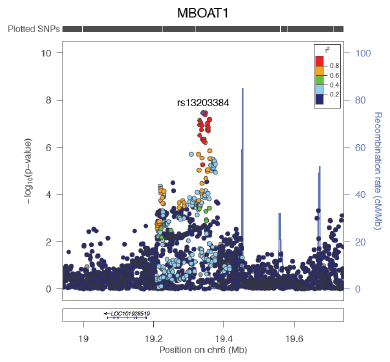

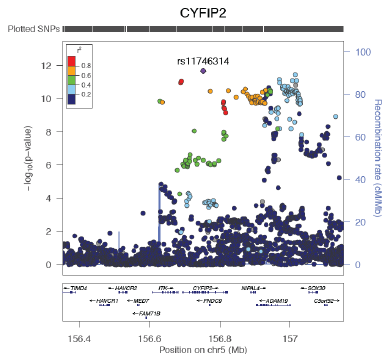

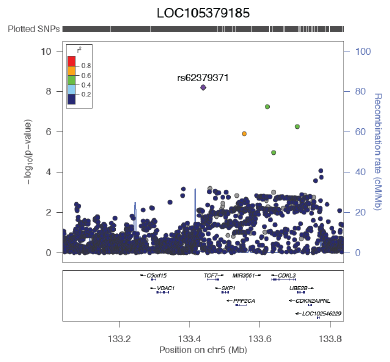

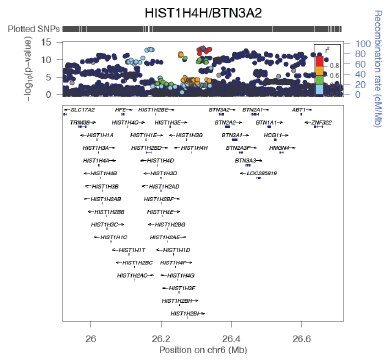

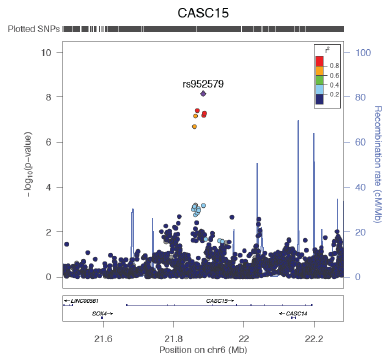

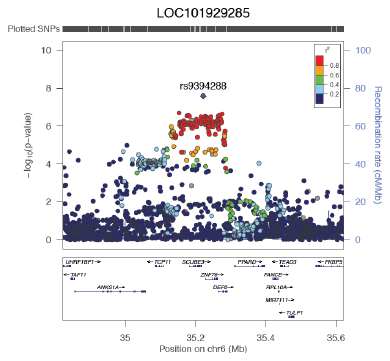

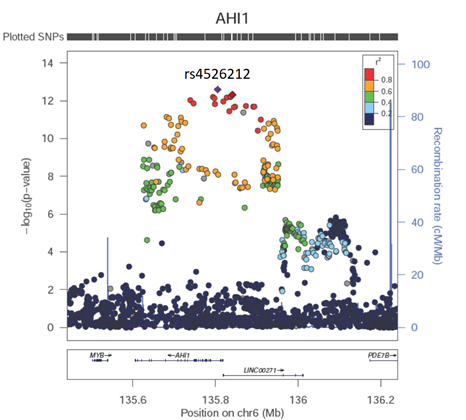

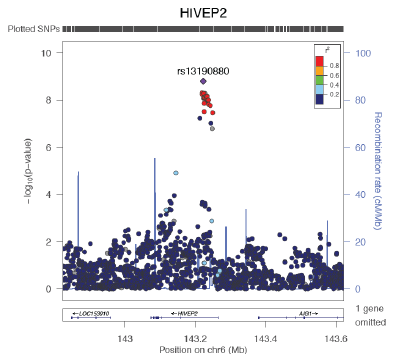

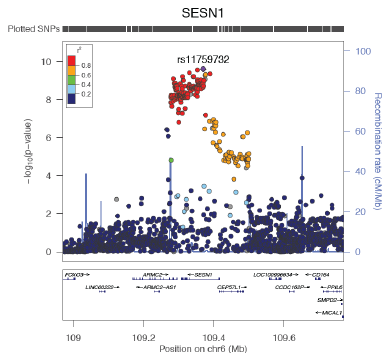

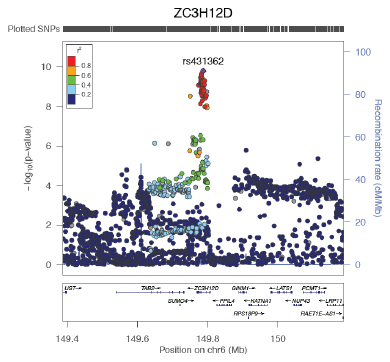

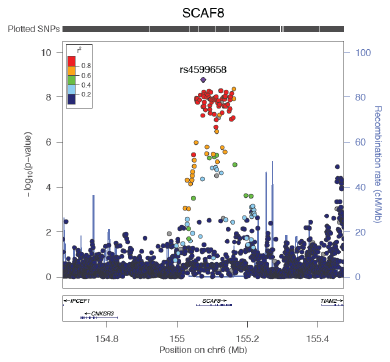

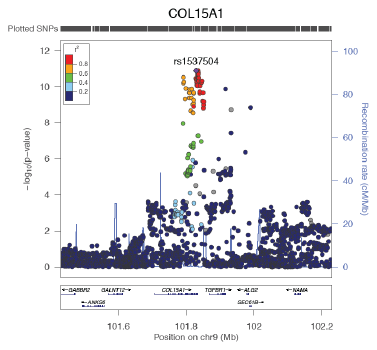

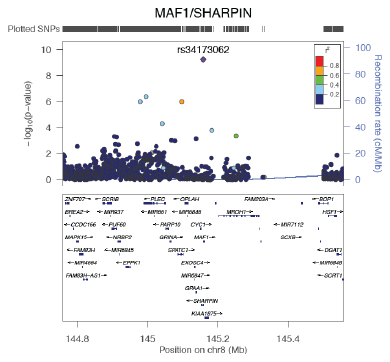

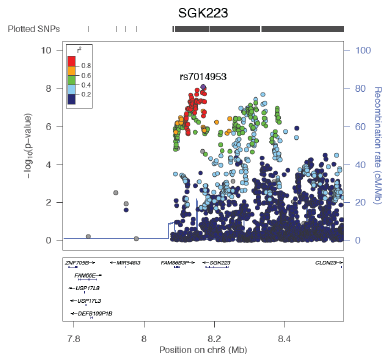

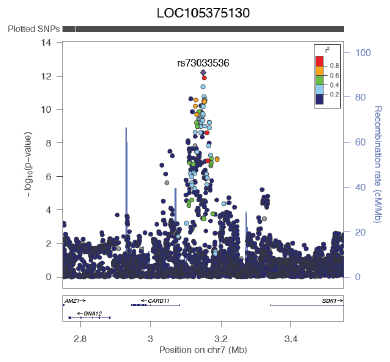

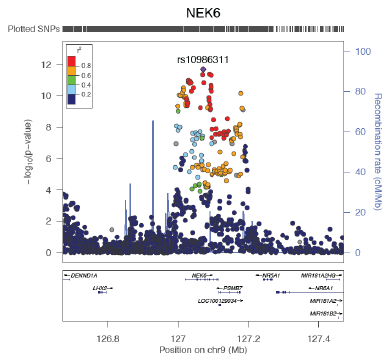

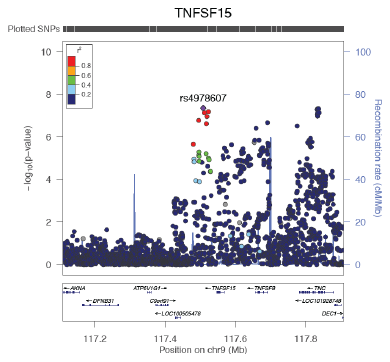

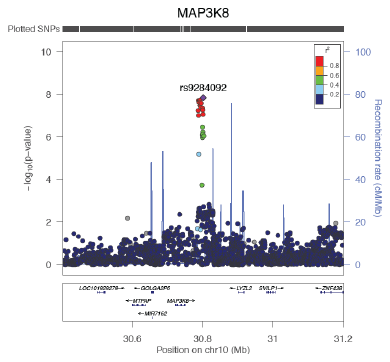

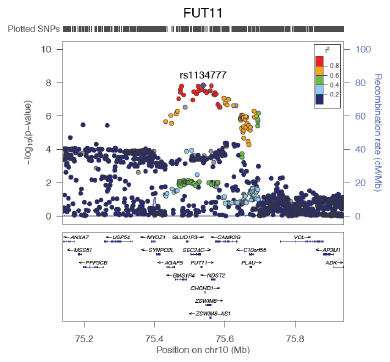

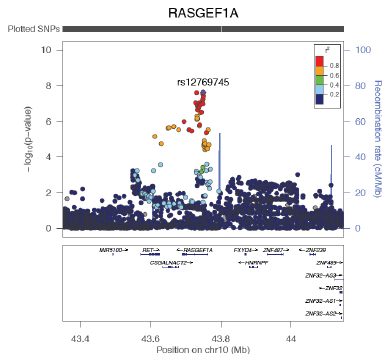

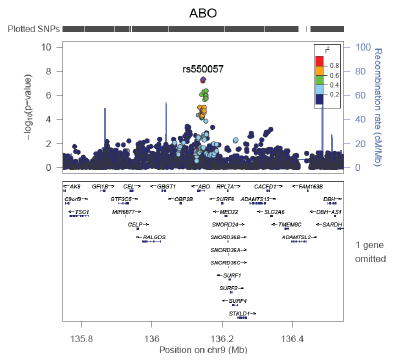

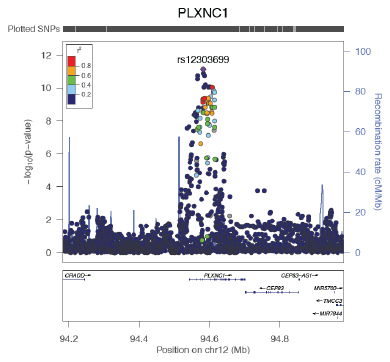

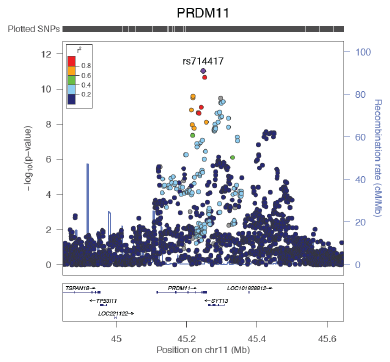

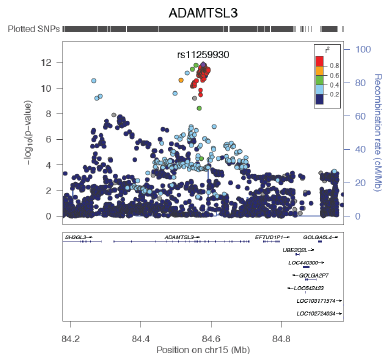

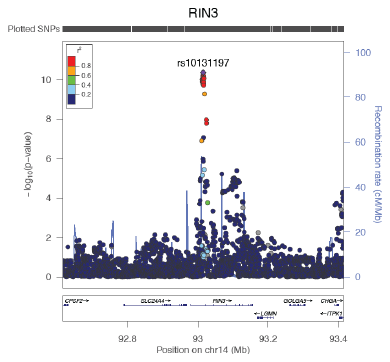

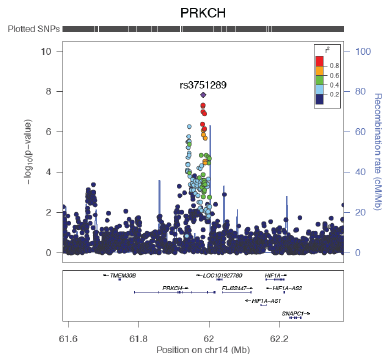

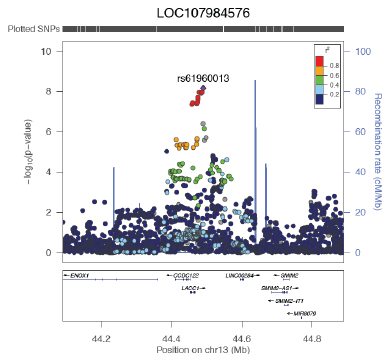

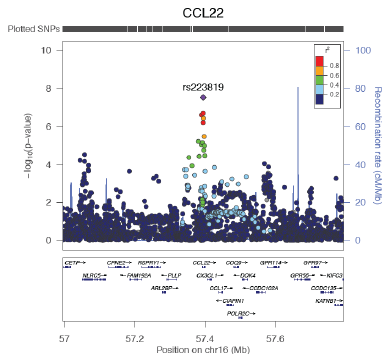

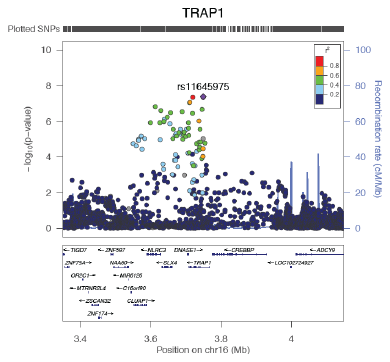

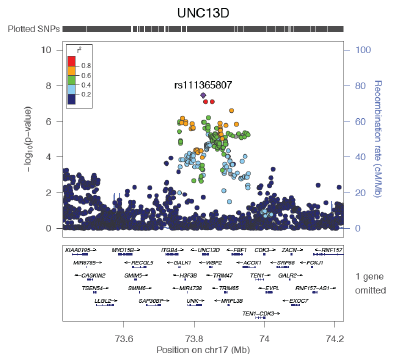

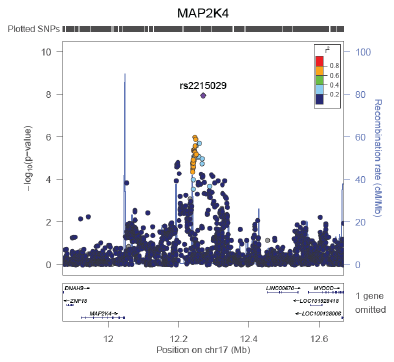

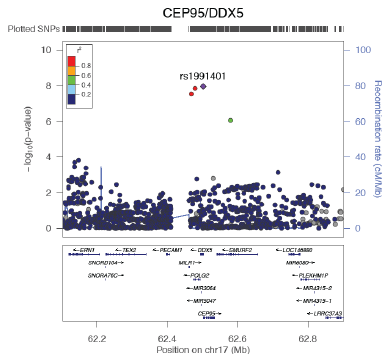

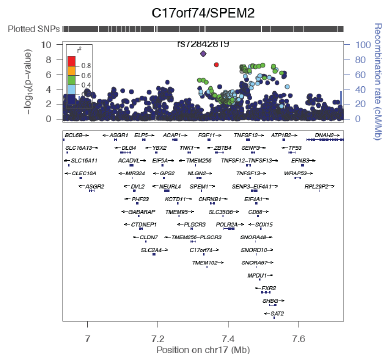

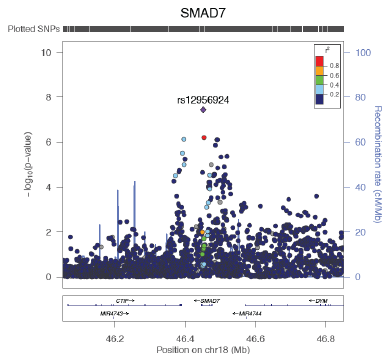

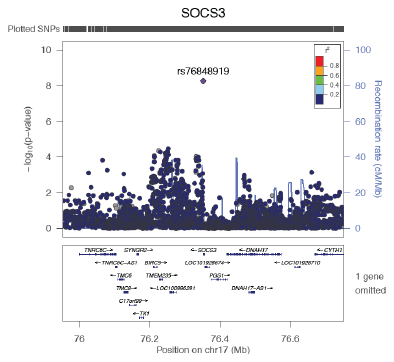

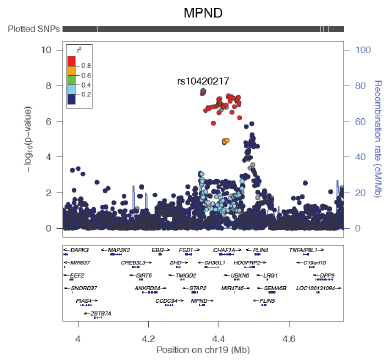

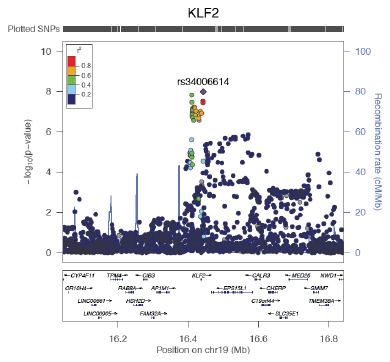

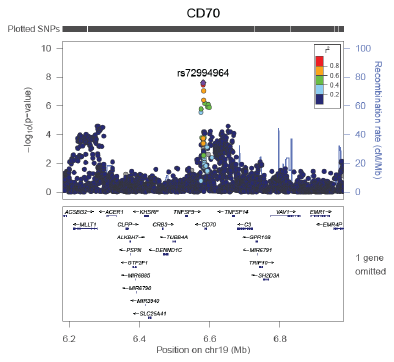

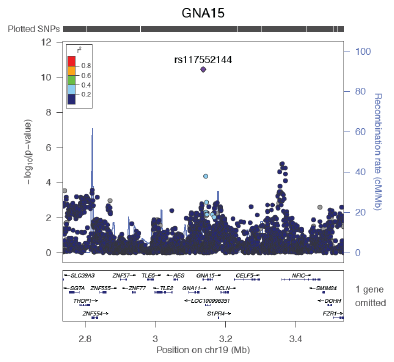

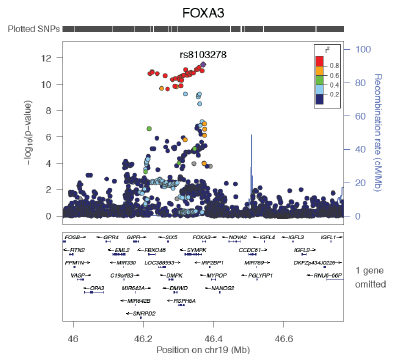

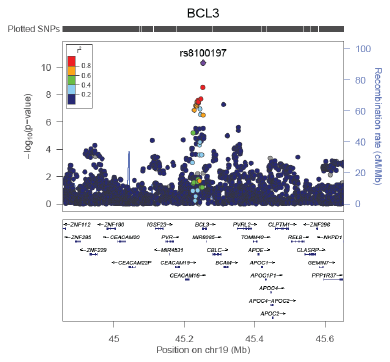

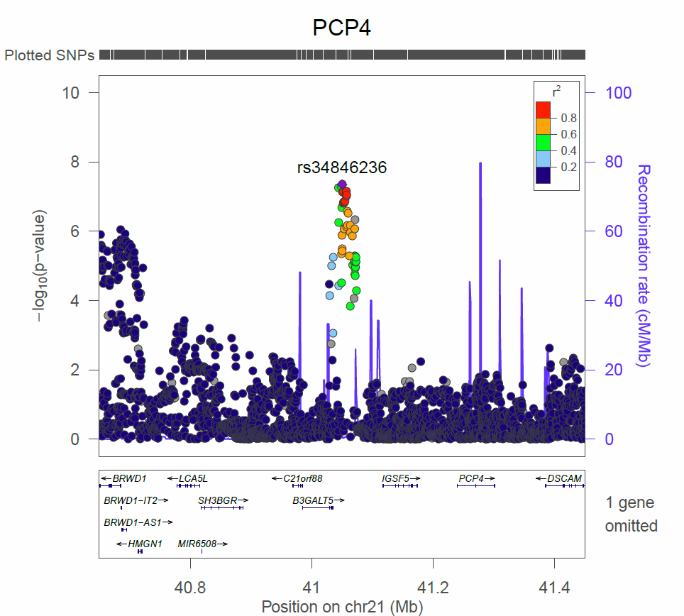

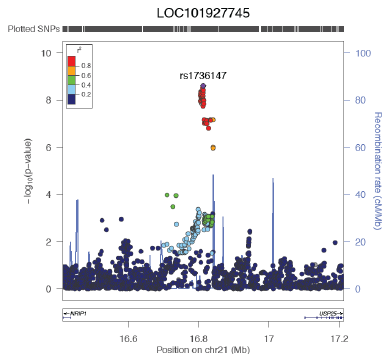

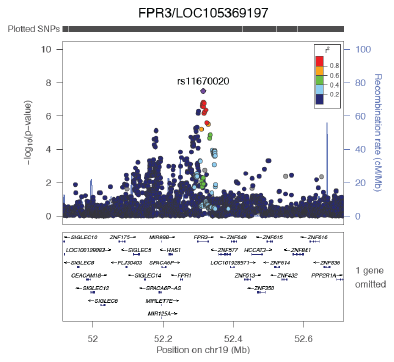

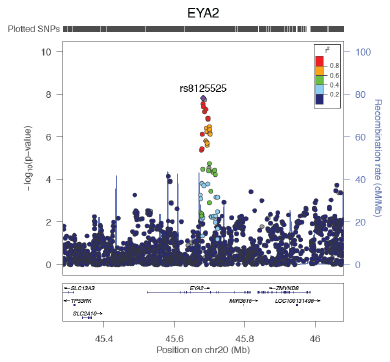

**Supplementary Figure 3. Regional plots of 66 novel loci identified for asthma in GWAS and meta-analyses with UK Biobank and TAGC.** 58 loci were identified as being significantly associated with asthma in the meta-analysis of UK Biobank and TAGC, whereas 8 loci (*RAF1*, *LOC105379185*, *LOC107984576*, *CEP95*-*DDX5*, *SOCS3*, *SMAD7*, *GNA15*, *MPND*) were genome-wide significant in the GWAS for asthma in UK Biobank alone. Each region is centered on the lead SNP (purple diamond) and the genes in the interval are indicated in the bottom panel. The degree of linkage disequilibrium (LD) between the lead SNP and other variants is shown as r^2^ values according to the color-coded legend in the box.

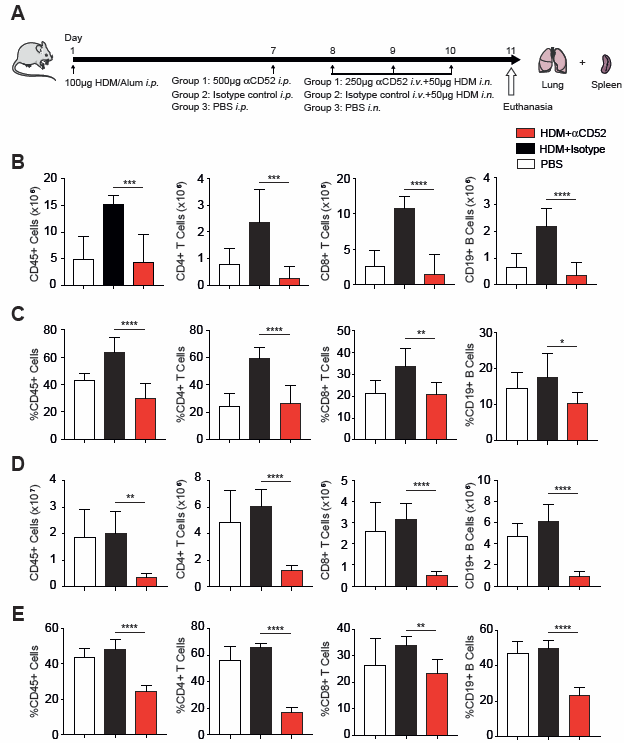

**Supplementary Figure 4. Mouse anti-CD52 (αCD52) antibody depletes lymphocytes in lung and spleen.** (**A)** Female BALB/cByJ mice were immunized on day 1 with 100µg of house dust mite (HDM) in 2mg of aluminum hydroxide (alum) by intraperitoneal (*i.p.*) injection. On day 7, mice were intraperitoneally administered with either 500µg of αCD52 antibody (Group 1; n=8), 500µg isotype control antibody (Group 2; n=9), or PBS (Group 3; n=4). On days 8, 9 and 10, mice in Groups 1 and 2 were intravenously (*i.v.*) administered 250µg αCD52 antibody or isotype control antibody, respectively, and simultaneously challenged intranasally (*i.n.*) with 50µg HDM. Mice in Group 3 were only challenged intranasally with PBS. On day 11, mice were euthanized and relative and absolute numbers of lymphocytes in lung and spleen were quantified by means of flow cytometry (see Methods for details). The number **(B)** and percentage **(C)** of CD45^+^ cells, CD4^+^ and CD8^+^ T cells, and CD19^+^ B cells were significantly lower in lungs of mice treated with αCD52 antibody and exposed to HDM compared to the HDM-exposed isotype antibody control group. **(D)** and **(E)** Treatment of HDM-exposed mice with αCD52 antibody similarly depleted splenic CD45^+^ lymphocytes, T cells, and B cells. Data are shown as mean ± SE. *P<0.05, **P<0.005; ***P<0.0005; ****P<0.0001.

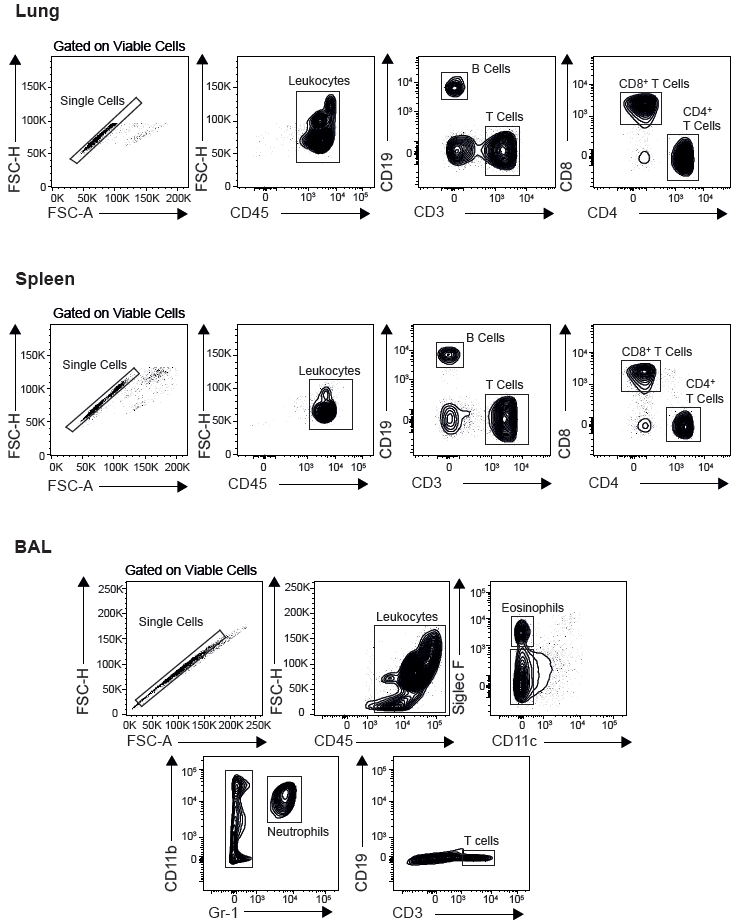

**Supplementary Figure 5. Gating strategy used to quantitate immune cells by flow cytometry.**  Representative flow plots showing that bone marrow-derived lymphocytes in lung and spleen were identified by gating on CD45, followed by gating on CD19 and CD3 for B and T cells, respectively. T cells were further gated on CD4 and CD8 to quantitate these subpopulations. A similar strategy was used in bronchial alveolar lavage to quantitate bone marrow-derived lymphocytes by gating on CD45, followed by gating on Siglec-F and CD11c for

eosinophils, Gr-1 and CD11b for neutrophils, and CD3 for T cells.
